# SARS-CoV-2 B.1.617.2 Delta variant replication, sensitivity to neutralising antibodies and vaccine breakthrough

**DOI:** 10.1101/2021.05.08.443253

**Authors:** Petra Mlcochova, Steven Kemp, Mahesh Shanker Dhar, Guido Papa, Bo Meng, Swapnil Mishra, Charlie Whittaker, Thomas Mellan, Isabella Ferreira, Rawlings Datir, Dami A. Collier, Anna Albecka, Sujeet Singh, Rajesh Pandey, Jonathan Brown, Jie Zhou, Niluka Goonawardne, Robin Marwal, Meena Datta, Shantanu Sengupta, Kalaiarasan Ponnusamy, Venkatraman Srinivasan Radhakrishnan, Adam Abdullahi, Oscar Charles, Partha Chattopadhyay, Priti Devi, Daniela Caputo, Tom Peacock, Chand Wattal, Neeraj Goel, Ambrish Satwik, Raju Vaishya, Meenakshi Agarwal, The Indian SARS-CoV-2 Genomics Consortium (INSACOG), The CITIID-NIHR BioResource COVID-19 Collaboration, The Genotype to Phenotype Japan (G2P-Japan) Consortium, Antranik Mavousian, Joo Hyeon Lee, Jessica Bassi, Chiara Silacci-Fegni, Christian Saliba, Dora Pinto, Takashi Irie, Isao Yoshida, William L. Hamilton, Kei Sato, Leo James, Davide Corti, Luca Piccoli, Samir Bhatt, Seth Flaxman, Wendy S. Barclay, Partha Rakshit, Anurag Agrawal, Ravindra K. Gupta

**Author notes:** Authors contributed equally to this work (Petra Mlcochova, Steven Kemp, Mahesh Shanker Dhar, Partha Rakshit, Anurag Agrawal, Ravindra K. Gupta).

## Abstract

The SARS-CoV-2 B.1.617.2 (Delta) variant was first identified in the state of Maharashtra in late 2020 and spread throughout India, outcompeting pre-existing lineages including B.1.617.1 (Kappa) and B.1.1.7 (Alpha). *In vitro*, B.1.617.2 is 6-fold less sensitive to serum neutralising antibodies from recovered individuals, and 8-fold less sensitive to vaccine-elicited antibodies as compared to wild type Wuhan-1 bearing D614G. Serum neutralising titres against B.1.617.2 were lower in ChAdOx-1 versus BNT162b2 vaccinees. B.1.617.2 spike pseudotyped viruses exhibited compromised sensitivity to monoclonal antibodies against the receptor binding domain (RBD) and N-terminal domain (NTD), in particular to the clinically approved bamlavinimab and imdevimab monoclonal antibodies. B.1.617.2 demonstrated higher replication efficiency in both airway organoid and human airway epithelial systems as compared to B.1.1.7, associated with B.1.617.2 spike being in a predominantly cleaved state compared to B.1.1.7. Additionally we observed that B.1.617.2 had higher replication and spike mediated entry as compared to B.1.617.1, potentially explaining B.1.617.2 dominance. In an analysis of over 130 SARS-CoV-2 infected healthcare workers across three centres in India during a period of mixed lineage circulation, we observed substantially reduced ChAdOx-1 vaccine efficacy against B.1.617.2 relative to non-B.1.617.2. Compromised vaccine efficacy against the highly fit and immune evasive B.1.617.2 Delta variant warrants continued infection control measures in the post-vaccination era.

## Introduction

India’s first wave of SARS-CoV-2 infections in mid-2020 was relatively mild and was controlled by a nationwide lockdown. Since easing of restrictions, India has seen expansion in cases of COVID-19 since March 2021 with widespread fatalities and a death toll of over 400,000. The B.1.1.7 Alpha variant, introduced by travel from the United Kingdom (UK) in late 2020, expanded in the north of India and is known to be more transmissible than previous viruses bearing the D614G spike mutation, whilst maintaining sensitivity to vaccine elicited neutralising antibodies^1,2^. The B.1.617 variant was first identified in the state of Maharashtra in late 2020/early 2021^3^, spreading throughout India and to at least 90 countries.

The first sub-lineage to be detected was B. 1.617.1^4–6^, followed by B. 1.617.2, both bearing the L452R spike receptor binding motif mutation also observed in B.1.427/B. 1.429^7,8^. This mutation was previously reported to confer increased infectivity and a modest loss of susceptibility to neutralising antibodies^9,10^. B.1.617.2, termed the Delta variant by WHO, has since dominated over B.1.617.1 (Kappa variant) and other lineages including B.1.1.7 globally (https://nextstrain.org/sars-cov-2)^11^. B.1.617.2 bears spike mutations T19R, G142D, E156G, F157del, R158del, L452R, T478K, D614G, P681R and D950N relative to Wuhan-1 D614G.

Although vaccines have been available since early 2021, achieving near universal coverage in adults has been an immense logistical challenge, in particular for populous nations where B.1.617.2 is growing rapidly with considerable morbidity and mortality^12^. Current vaccines were designed to target the B.1, Wuhan-1 virus, and the emergence of variants with reduced susceptibility to vaccines such as B.1.351 and P.1 has raised fears for longer term control and protection through vaccination^13,14^, particularly in risk groups^15,16^. The specific reasons behind the explosive global growth of B.1.617.2 in populations remain unclear. Possible explanations include evasion of neutralising antibodies generated through vaccination or prior infection, as well as increased infectivity.

## Results

### SARS-CoV-2 B.1.617.2 shows reduced sensitivity to neutralising antibodies

We first plotted the relative proportion of variants in new cases of SARS-CoV-2 in India since the start of 2021. Whilst B.1.617.1 emerged earlier, it has been replaced by the Delta variant B.1.617.2 (**Figure 1a**). We hypothesised that B.1.617.2 would exhibit immune evasion to antibody responses generated by previous SARS-CoV-2 infection. We used sera from twelve individuals infected during the first UK wave in mid-2020 (likely following infection with SARS-CoV-2 Wuhan-1). These sera were tested for ability to neutralise a B.1.617.2 viral isolate (obtained from nose/throat swab), in comparison to a B.1.1.7 variant isolate and a wild type (WT) Wuhan-1 virus bearing D614G in spike. The Delta variant contains several spike mutations that are located at positions within the structure that are predicted to alter its function (**Figure 1b**). We found that the B.1.1.7 virus isolate was 2.3-fold less sensitive to the sera compared to the WT, and that B.1.617.2 was 5.7-fold less sensitive to the sera (**Figure 1c**). Importantly in the same assay, the B.1.351 Beta variant that emerged in South Africa demonstrated an 8.2-fold loss of neutralisation sensitivity relative to WT.

**Figure. 1:**
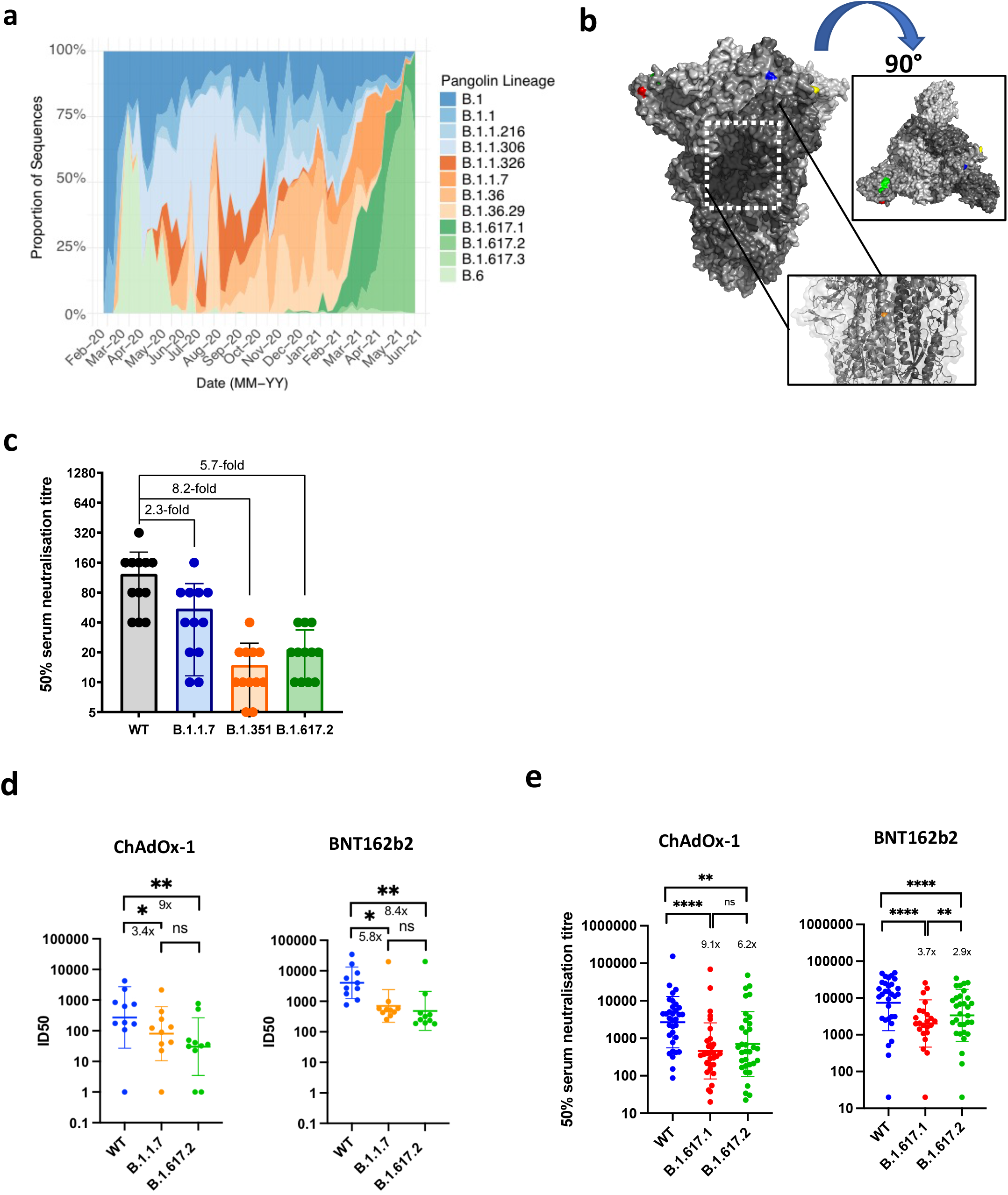
Rapid Expansion of Delta variant B.1.617.2 in India and reduced sensitivity to neutralizing antibodies from sera derived following infection and vaccination. **a.** Proportion of lineages in incident cases of SARS-CoV-2 in India 2020-2021. **b.** Surface representation of the SARS-CoV-2 B.1.671.2 Spike trimer (PDB: 6ZGE). L19R (red), del157/158 (green) L452R (blue) and T478K (yellow). The white dashed box indicates the location of the D950N (orange) **c. Neutralization of Delta variant by convalescent human serum from mid-2020** in Vero-hACE2 TMPRSS2 cells. Fold-change in serum neutralisation 100 TCID_50_ of B.1.17 (Alpha), B.1.351 (Beta) and B.1617.2 (Delta) variants relative to wild-type (IC19), n=12. **d.** Neutralisation of delta variant live virus isolate by sera from vaccinated individuals (n=10 ChAdOx-1 or n=10 BNT12b2) in comparison to B.1.1.7 and Wuhan-1 wild type (WT). **5**-fold dilutions of vaccinee sera were mixed with wild type (WT) or viral variants (MOI 0.1) for 1h at 37°C. Mixture was added to Vero-hACE2/TMPRSS2 cells for 72h. Cells were fixed and stained with Coomasie blue and % of survival calculated. ID50 were calculated using nonlinear regression. Graph represents average of two independent experiments. **e.** Neutralisation of B.1.617 spike pseudotyped virus (PV) and wild type (WT, Wu-1 D614G) by vaccine sera (n=33 ChAdOx-1 or n=32 BNT162b2). GMT (geometric mean titre) with s.d are presented. Data representative of two independent experiments each with two technical replicates. **p<0.01, *** p<0.001, ****p<0.0001 Wilcoxon matched-pairs signed rank test, ns not significant.

We used the same B.1.617.2 live virus isolate to test susceptibility to vaccine elicited serum neutralising antibodies in individuals following vaccination with two doses ChAdOx-1 or BNT162b2. These experiments showed a loss of sensitivity for B.1.617.2 compared to wild type Wuhan-1 bearing D614G of around 8-fold for both sets of vaccine sera and reduction against B.1.1.7 that did not reach statistical significance (**Figure 1d**). We also used a pseudotyped virus (PV) system to test neutralisation potency of a larger panel of 65 vaccine-elicited sera, this time against B.1.617.1 as well as B.1.617.2 spike compared to Wuhan-1 D614G spike (**Figure 1e**). Comparison of demographic data for vaccinees showed similar characteristics (**Extended Data Table 1**). The mean GMT against Delta Variant spike PV was lower for ChAdOx-1 compared to BNT162b2 (GMT 3372 versus 654, p<0001, **Extended Data Table 1**).

We investigated the role of the B.1.617.2 spike as an escape mechanism by testing 33 (3 NTD, 21 RBM- and 9 non-RBM-specific) spike-specific mAbs isolated from 6 individuals that recovered from WT SARS-CoV-2 infection with an *in-vitro* PV neutralization assay using Vero E6 target cells expressing Transmembrane protease serine 2 (TMPRSS2) and the Wuhan-1 D614G SARS-CoV-2 spike or the B.1.617.2 spike (**Figure 2a-c**, **Extended Data Figure 1a-c and Extended Data Table 2**). In addition, 5 clinical-stage RBM-mAbs (etesevimab, casirivimab, regdanvimab, imdevimab and bamlanivimab) were also tested using Vero E6 cells (**Figure 2c, Extended Data Figure 1d and Extended Data Table 2**). We found that all three NTD-mAbs (100%) and four out of nine (44%) non-RBM mAbs completely lost neutralizing activity against B.1.617.2 (**Figure 2 b-c and Extended Data Figure 1a**). Within the RBM-binding group, 16 out 26 mAbs (61.5%) showed a marked decrease (2-35 fold-change reduction) or complete loss (>40 fold-change reduction) of neutralizing activity to B.1.617.2, suggesting that in a sizeable fraction of RBM antibodies the L452R and T478K mutations are responsible for their loss of neutralizing activity (**Figures 2b-c, Extended Data Figure 1b**). Amongst the clinical-stage RBM-mAbs tested, bamlanivimab, which showed benefit in a clinical trial against prior variants^17^, did not neutralize B.1.617.2. Imdevimab, part of the REGN-COV2 therapeutic dual antibody cocktail^18^, displayed reduced neutralizing activity in Vero E6-TMPRSS2 cells (**Figure 2c and Extended Data Figure 1d-f**). The remaining clinical-stage mAbs, including S309 (the parental antibody from which sotrovimab was derived), retained potent neutralizing activity against B.1.617.2.

**Figure. 2:**
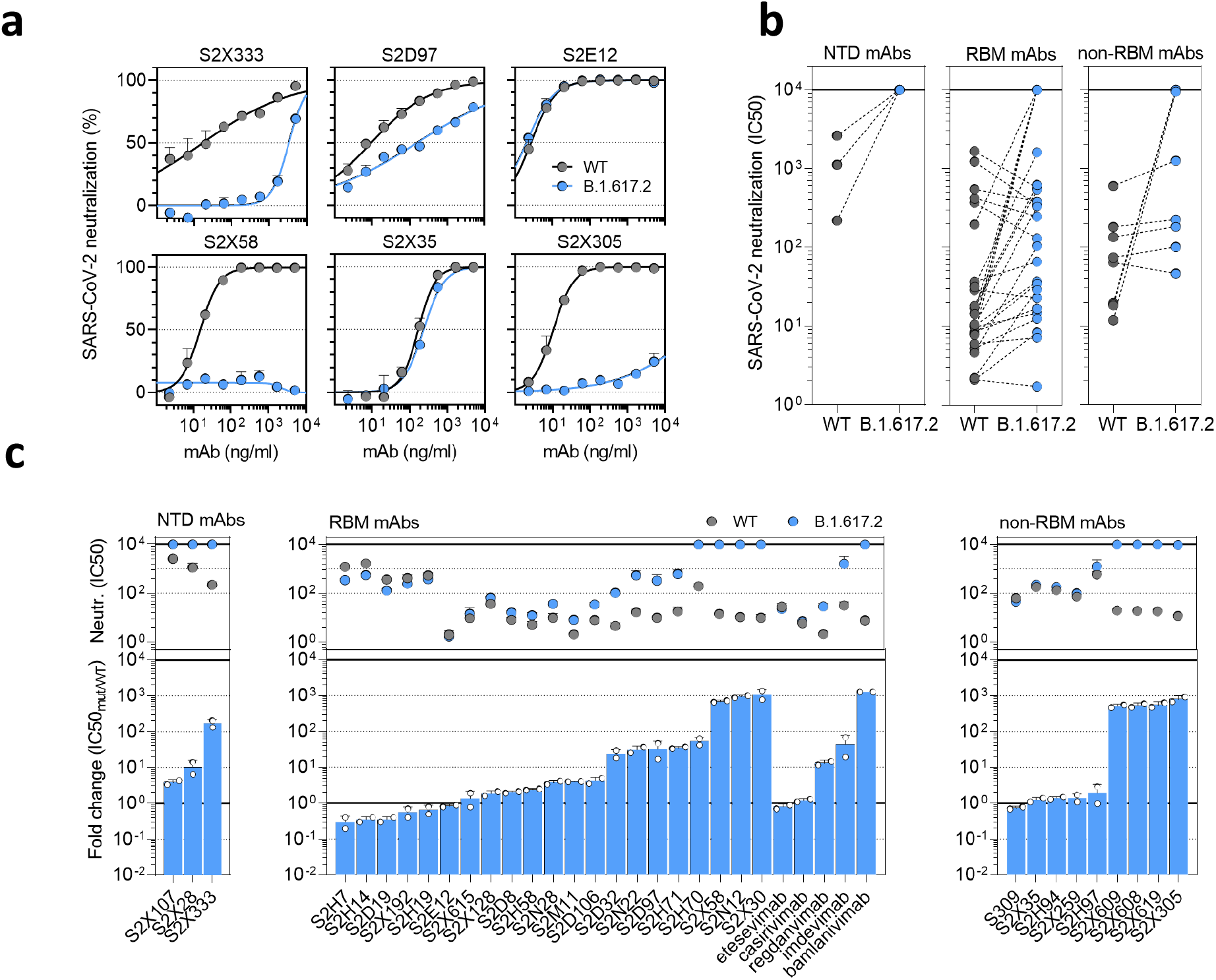
Delta variant B.1.617.2 shows reduced sensitivity to monoclonal antibodies. **a.** Neutralisation of WT D614 (black) and B.1.617.2 mutant (blue) pseudotyped SARS-CoV-2-VSV by 6 selected mAbs from one representative experiment out of 2 independent experiments. S2X333 is an NTD-specific mAb, S2D97, S2E12 and S2X58 are RBM-specific mAbs, while S2X35 and S2X305 are non-RBM mAbs. **b.** Neutralisation of WT and B.1.617.2 VSV by 38 mAbs targeting NTD (n=3), RBM (n=26, including 5 clinical stage mAb) and non-RBM (n=9). Shown are the mean IC50 values (ng/ml) from 2 independent experiments. Non-neutralising IC50 titers were set at 10^4^ ng/ml. **c**. Neutralisation shown as mean IC50 values (upper panel) and average fold change of B.1.617.2 relative to WT (lower panel) of 38 mAbs tested in 2 independent experiments (including 5 clinical-stage mAbs), tested using Vero E6 cells expressing TMPRSS2.

### SARS-CoV-2 B.1.617.2 variant shows higher replication in human airway model systems

We next sought biological evidence for the higher transmissibility predicted from the modelling. Increased replication could be responsible for generating greater numbers of virus particles, or the particles themselves could be more likely to lead to a productive infection. We first infected a lung epithelial cell line, Calu-3, comparing B.1.1.7 and B.1.617.2 (**Figure 3a-d**). We observed a replication advantage for B.1.617.2 as demonstrated by intracellular RNA transcripts and S and N proteins (**Figure 3a-b**), as well as analysis of released virions from cells (**Figure 3c-d**). Next we tested B.1.1.7 against two separate isolates of B.1.617.2 in a human airway epithelial model^19^. In this system we again observed that both B.1.617.2 isolates had a significant replication advantage over B.1.1.7 (**Figure 3e-f**). Finally, we infected primary 3D airway organoids^20^ (**Figure 3g**) with B.1.617.2 and B.1.1.7 virus isolates, noting a significant replication advantage for B.1.617.2 over B.1.1.7. These data clearly support higher replication rate and therefore transmissibility of B.1.617.2 over B.1.1.7.

**Figure 3.**
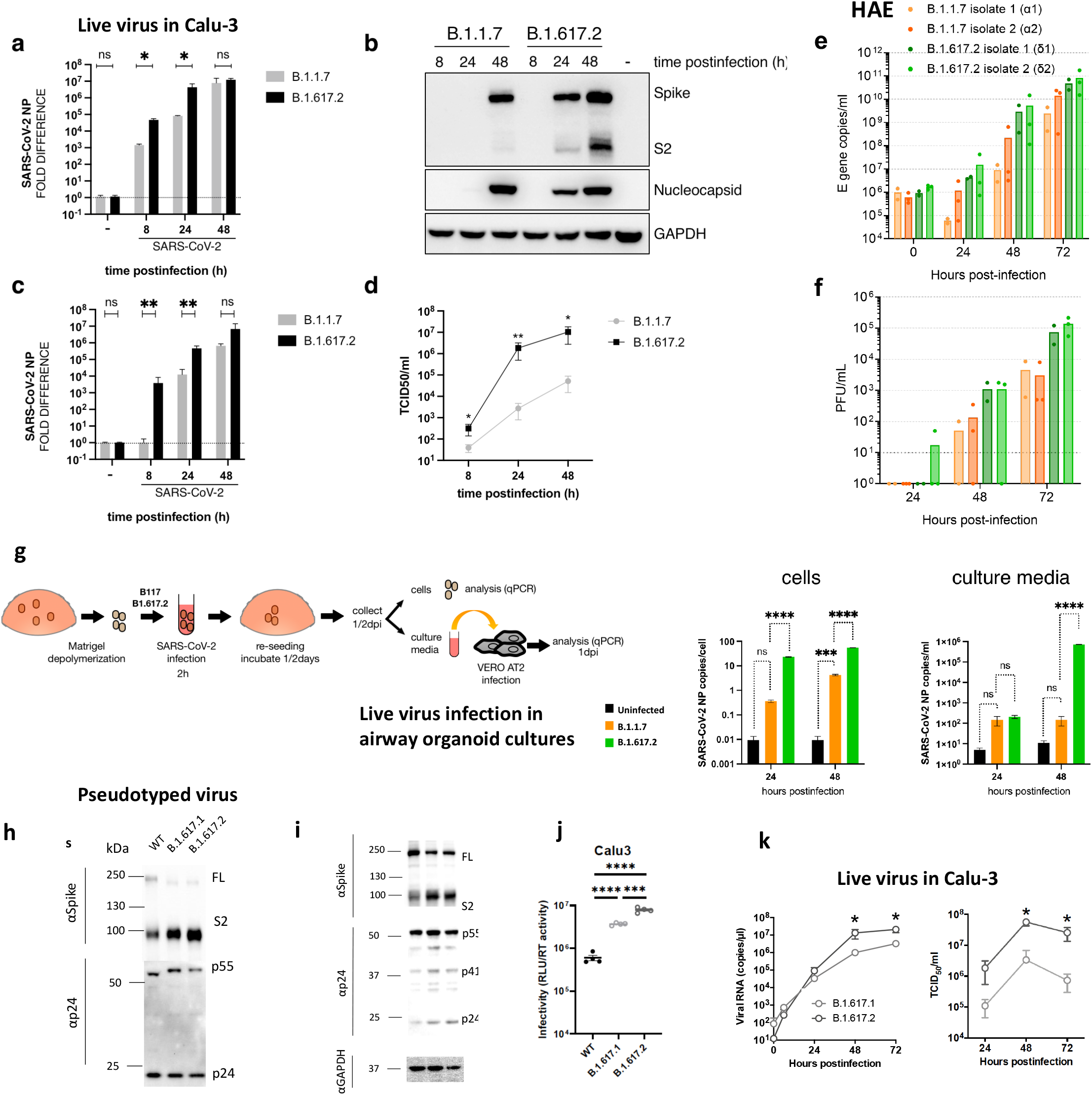
a. SARS-CoV-2 B.1.617.2 Delta Variant replication and and spike mediated entry efficiency. **a-d**. Live virus replication in CaLu-3 lung cells comparing B.1.1.7 with B.1.617.2. Calu-3 cells were; infected with variants at MOI 0.1. Cells and supernatants containing released virus were collected for RNA isolation, western blot and TCID_50_ at 8, 24 and 48h postinfection. **a.** viral loads were measured by qPCR in cell lysates. **b.** viral protein levels were detected in cell lysates. **c-d** Live virus produced from infected Calu3 cells was collected and used to infect permissive Vero Ace2/TMPRSS2 cells to measure **c**. viral loads in Vero cells or **d**. to measure TCID_50_/ml. **e-f.** Virus replication kinetics in human airway epithelial (HAE) system with air liquid interface ALI, using two B.1.617.2 isolates and two B.1.1.7 isolates. **g.** Live virus **r**eplication in airway epithelial organoid cultures. Airway epithelial organoids were infected with SARS-CoV-2 variants B.1.1.7 and B.1.617.2 at MOI 1. Cells were lysed 24 and 48h post-infection and total RNA isolated. qPCR was used to determine copies of nucleoprotein gene in organoid cells and infectivity of cell free virus measured by infection of Vero AT2 cells. Data represent are representative of two independent experiments, **h** and **i.** western blots of pseudotyped virus (PV) virions (h) and cell lysates (i) of 293T producer cells following transfection with plasmids expressing lentiviral vectors and SARS-CoV-2 S B.1.617.1 and Delta variant B.1.617.2 versus WT (Wuhan-1 with D614G), probed with antibodies for HIV-1 p24 and SARS-Cov-2 S2. **j.** Single round infectivity on Calu-3 by spike B.1.617.2 and B.1.617.1 versus WT D614G parental plasmid PV produced in 293T cells. Data are representative of three independent experiments. **k.** Growth kinetics of B.1.617.1 and B.1.617.2 variants. Viral isolates of B.1.617.1 and B.1.617.2 [200 50% tissue culture infectious dose (TCID_50_)] were inoculated into Calu-3 cells and the copy number of viral RNA in the culture supernatant was quantified by real-time RT-PCR over time. TCID_50_ of released virus in supernatant was also measured over time. Assays were performed in quadruplicate. *, *P*<0.05 by Mann-Whitney U test.. ns, non-significant; * p<0.05; ** p < 0.01. ***p<0.001, ****p<0.0001 (-) uninfected cells. Data are representative of two independent experiments

In the aforementioned experiments we noted a higher proportion of intracellular B.1.617.2 spike in the cleaved state in comparison to B.1.1.7 (**Figure 3b**). In order to investigate this further we produced the two viruses as well as a B.1 D614G virus in Vero-hACE2-TMPRSS2 cells, harvested and purified supernatants at 48 hours before running western blots probing for spike S2 and nucleoprotein. This analysis showed that the B.1.617.2 spike was predominantly in the cleaved form, in contrast to B.1 and B.1.1.7 (**Extended Data Figure 2a-b**).

### SARS-CoV-2 B.1.617.2 spike has enhanced entry efficiency associated with cleaved spike

SARS-CoV-2 Spike is known to mediate cell entry via interaction with ACE2 and TMPRSS2^21^ and is a major determinant of viral infectivity. In order to gain insight into the mechanism of increased infectivity of B.1.617.2, we tested single round viral entry of B.1.617.1 and B.1.617.2 spikes (**Figure 3h,i** and **Extended Data Figure 3a-b**) using the pseudotyped virus (PV) system, infecting Calu-3 lung cells expressing endogenous levels of ACE2 (Angiotensin Converting Enzyme 2) and TMPRSS2 (Transmembrane protease serine 2) (**Figure 3j**), as well as other cells transduced or transiently transfected with ACE2 / TMPRSS2 (**Extended Data Figure 3b**). We first probed PV virions and cell lysates for spike protein and noted that the B.1.617 spikes were present predominantly in cleaved form in cells and virions, in contrast to WT (**Figure 3h-i, Extended Data Figure 3c**). We observed one log increased entry efficiency for both B.1.617.1 and B.1.617.2. over Wuhan-1 D614G wild type in nearly all cells tested (**Extended Data Figure 3b**). In addition, B.1.617.2 appeared to have an entry advantage compared to B.1.617.1 in some cells, and in particular Calu-3 bearing endogenous receptors (**Figure 3j**). Finally, we wished to confirm higher infectivity using live virus isolates of B.1.617.1 and B.1.617.2. As expected from the PV comparison, B.1.617.2 showed increased replication kinetics B.1.617.1 in Calu-3 cells over 48 hours as measured by supernatant RNA and TCID_50_ (**Figure 3k**).

### SARS-CoV-2 B.1.617.2 spike confers increased syncytium formation

The plasma membrane route of entry, and indeed transmissibility in animal models, is critically dependent on the polybasic cleavage site (PBCS) between S1 and S2^19,22,23^ and cleavage of spike prior to virion release from producer cells; this contrasts with the endosomal entry route, which does not require spike cleavage in producer cells.^19,24,25^. Mutations at P681 in the PBCS have been observed in multiple SARS-CoV-2 lineages, most notably in the B.1.1.7 Alpha variant^24^. We previously showed that B.1.1.7 spike, bearing P681H, had significantly higher fusogenic potential than a D614G Wuhan-1 virus^24^. Here we tested B.1.617.1 and B.1.617.2 spike using a split GFP system to monitor cell-cell fusion (**Figure 4a, b, c**). We transfected spike bearing plasmids into Vero cells stably expressing the two different parts of Split-GFP, so that GFP signal could be measured over time upon cell-cell fusion (**Figure 4d**). The B.1.617.1 and B.1.617.2 spike proteins mediated higher fusion activity and syncytium formation than WT, and were similar to B.1.1.7 (**Figure 4d,e**). The single P681R mutation was able to recapitulate this phenotype (**Figure 4d,e**). Finally we explored whether post vaccine sera could block syncytia formation, as this might be a mechanism for vaccine protection against pathogenesis. We titrated sera from ChAdOx-1 vaccinees and showed that indeed the cell-cell fusion could be inhibited in a manner that mirrored neutralisation activity of the sera against PV infection of cells (**Figure 4f**). Hence B.1.617.2 may induce cell-cell fusion in the respiratory tract and possibly higher pathogenicity even in vaccinated individuals with neutralising antibodies.

**Figure. 4:**
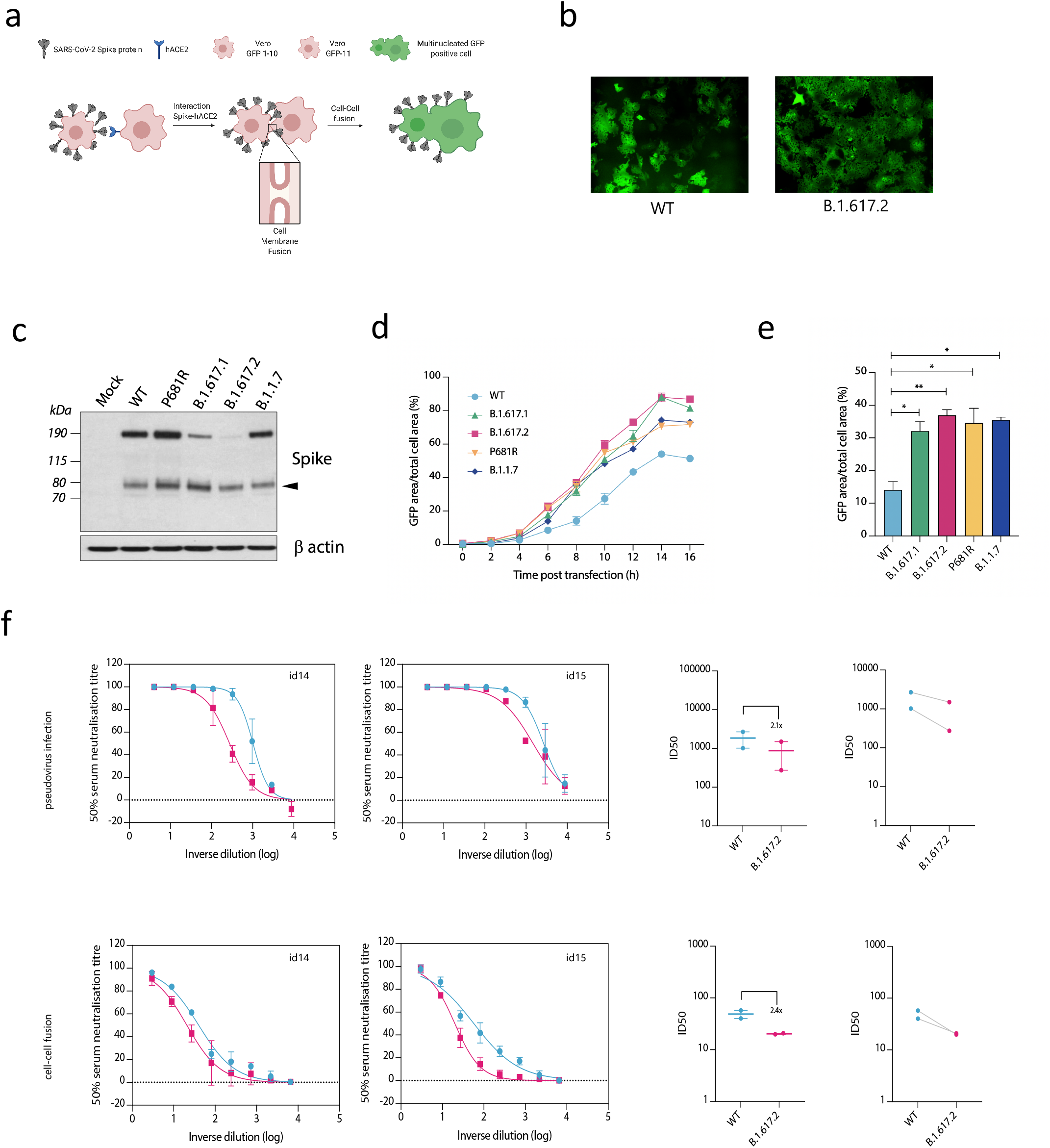
B.1.617.2 Delta variant spike confers accelerated cell-cell fusion activity that can be blocked by anti-spike neutralising antibodies in sera. **a.** Schematic of cell-cell fusion assay. **b.** Reconstructed images at 10 hours of GFP positive syncytia formation. Scale bars represent 400 mm. **c.** western blot of cell lysates 48 hours after transfection of spike plasmids. Anti-S2 antibody. **d,e.** Quantification of cell-cell fusion kinetics showing percentage of green area to total cell area over time. Mean is plotted with error bars representing SEM. **f.** Comparison of impact of post vaccine sera (n=2) on PV neutralisation (top) and cell-cell fusion (bottom), comparing WT and Delta variant B.1.671.2. Data are representative of at least two independent experiments.

### Breakthrough SARS-CoV-2 B.1.617.2 infections in vaccinated health care workers

Hitherto we have gathered epidemiological and biological evidence that the growth advantage of B.1.617.2 might relate to increased virus replication/transmissibility as well as re-infection due to evasion of neutralising antibodies from prior infection. We hypothesised that vaccine effectiveness against B.1.617.2 would be compromised relative to other circulating variants. Although overall national vaccination rates were low in India in the first quarter of 2021, vaccination of health care workers (HCW) started in early 2021 with the ChAdOx-1 vaccine (Covishield). During the wave of infections during March and April, an outbreak of symptomatic SARS-CoV-2 was confirmed in 30 vaccinated staff members amongst an overall workforce of 3800 at a single tertiary centre in Delhi by RT-PCR of nasopharyngeal swabs (age range 27-77 years). Genomic data from India suggested B.1.1.7 dominance overall (**Figure 1a**) and in the Delhi area during the first quarter of 2021 (**Figure 5a**), with growth of B.1.617 during March 2021. By April 2021, 385 out of 604 sequences reported to GISAID for Delhi were B.1.617.2. Short-read sequencing^26^ of symptomatic individuals in the HCW outbreak revealed the majority were B.1.617.2 with a range of other B lineage viruses including B.1.1.7 and B.1.617.1 (**Figure 5b**). There were no cases that required ventilation though one HCW received oxygen therapy. Phylogenetic analysis demonstrated a group of highly related, and in some cases, genetically indistinct sequences that were sampled within one or two days of each other (**Figure 5b**). These data are consistent with a single transmission from an infected individual, constituting an over dispersion or ‘super spreader’ event. We next looked in greater detail at the vaccination history of cases. Nearly all had received two doses at least 21 days previously, and median time since second dose was 27 days.

**Figure 5.**
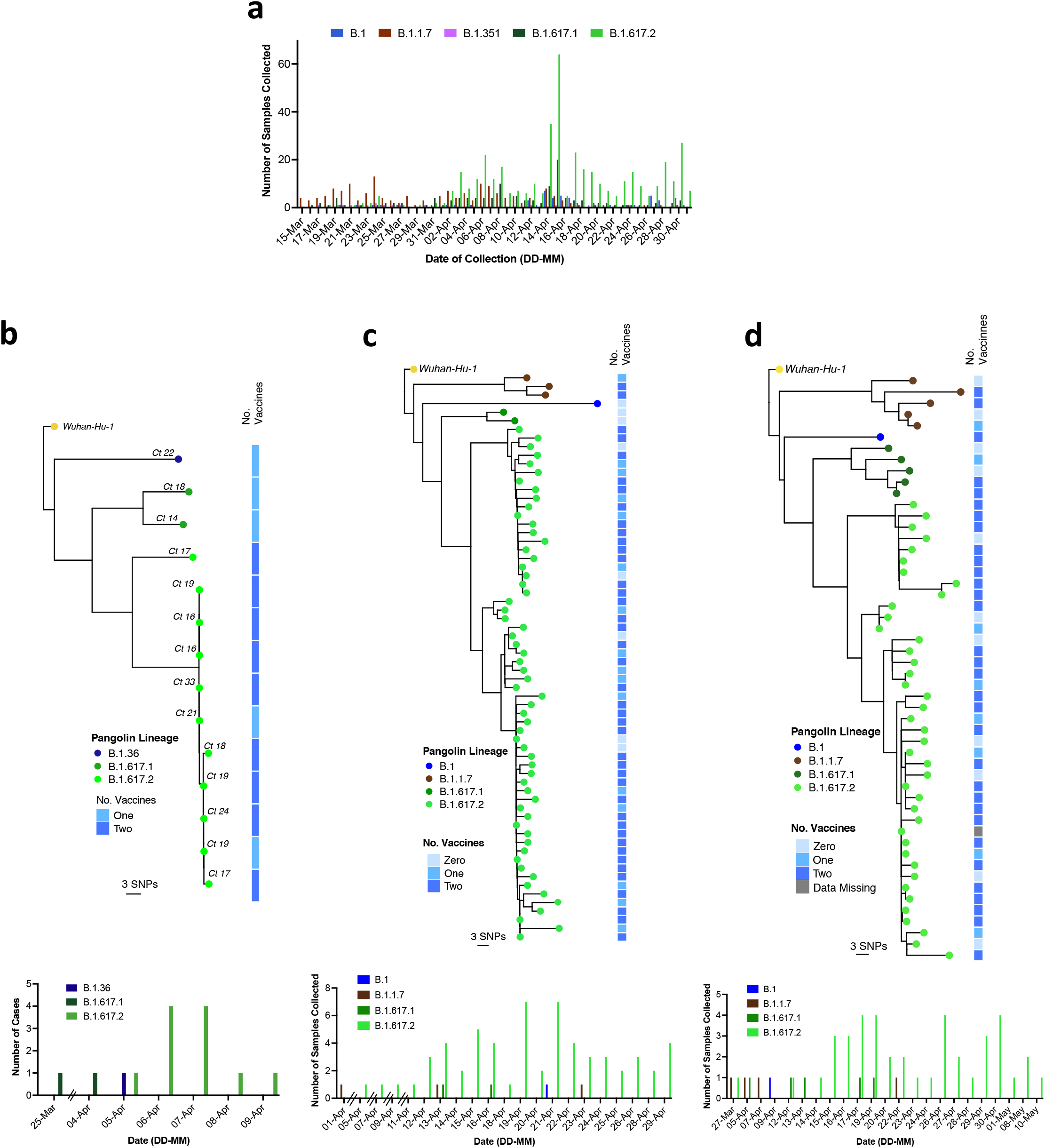
SARS-CoV-2 B.1.617.2 infection in vaccinated health care workers. **a.** Case frequencies of five most commonly occurring SARS CoV-2 lineages over a six week period from March to April 2021 for Delhi. **b-d** Maximum likelihood phylogenies of vaccine breakthrough SARS-CoV-2 sequences amongst vaccinated HCW at three centres are presented. Phylogenies were inferred with IQTREE2 with 1000 bootstrap replicates. Trees are rooted on Wuhan-Hu-1 and annotated with the lineage designated by pangolin v.2.4.2. The number of COVISHIELD (ChAdOx-1) vaccinations received by each individual is indicated by the heatmap to the right. White space indicates missing data. At the bottom of each tree is a case frequency graph by date of testing.

We obtained similar data on vaccine breakthrough infections in two other health facilities in Delhi with 1100 and 4000 HCW staff members respectively (**Figure 5c-d**). In hospital two there were 118 sequences from symptomatic, non-fatal infections, representing over 10% of the workforce over a 4 week period. After filtering, we reconstructed phylogenies using 66 with high quality whole genome coverage >95%. In hospital three there were 70 symptomatic, non-fatal infections from which genomes were generated, with 52 high quality genomes used for inferring phylogenies after filtering (**Figure 5c-d**). As expected from variants circulating in the community, we observed that B.1.617.2 dominated vaccine-breakthrough HCW infections (**Figure 5c-d**).

Across the three centres we noted that the median age of those infected with B.1.617.2 versus non-B.1.617.2 was similar [36.5 versus 32.5, p=0.56, (**Extended Data Table 3)**]. Half of breakthrough infections were in females regardless of variant. We observed no significant difference in the median duration of symptoms in B.1.617.2 versus non-B.1.617.2 infections (1.5 versus 1.0 days respectively, **Extended Data Table 3**), consistent with efficient symptomatic staff testing. Around 5% of symptomatic infections resulted in hospitalisation, with no evidence that B.1.617.2 was associated with higher risk of hospitalisation (**Extended Data Table 3**). The magnitude of vaccine responses in a limited sample of HCW with subsequent breakthrough was measured and appeared similar to responses in a control group of HCW that did not subsequently test positive for SARS-CoV-2 (**Extended Data Figure 4**). Analysis of Ct values in positive samples by hospital did not show significant differences between HCW infected with B.1.617.2 versus non-B.1.617.2 (**Extended Data Figure 4**).

Next, we evaluated the effect of B.1.617.2 on vaccine effectiveness (VE) against symptomatic infection in the HCWs as compared to other lineages. In terms of observational studies the test negative case control approach would be ideal. Given the lack of availability of test negative data in our HCW setting we used an alternative approach to estimate VE used by Public Health England (PHE)^27^. If the vaccine had equal effectiveness against B.1.617.2 and non-B.1.617.2, a similar proportion of B.1.617.2 and non-B.1.617.2 breakthrough cases would be expected in both vaccinated and unvaccinated individuals. However, in our HCW, non-B.1.617.2 was isolated in a lower proportion of symptomatic cases in the fully vaccinated group compared to unvaccinated cases (**Extended Data Table 4**). We used multivariable logistic regression to estimate the odds ratio of testing positive with B.1.617.2 versus non-B.1.617.2 in vaccinated relative to unvaccinated individuals, adjusting for age, sex and hospital. The adjusted odds ratio for B.1.617.2 relative to non-B.1.617.2 was 5.45 (95% CI 1.39-21.4, p=0.018) for two vaccine doses (**Extended Data Table 4**). Calendar time, often associated with vaccination status, was unlikely to be a significant confounder here given the short time period studied. The analysis presented, whilst limited by relatively small numbers of non-B.1.671.2 infections and potentially affected by unmeasured confounders, is nevertheless consistent with UK data where the non-B.1.617.2 infections were largely B.1.1.7^28^.

## Discussion

Here we have combined *in vitro* experimentation and molecular epidemiology to propose that increased replication fitness and reduced sensitivity of SARS-CoV-2 B.1.617.2 to neutralising antibodies have contributed to the recent rapid replacement of B.1.1.7 and other lineages by B.1.617.2 in countries such as India, the U.S and the U.K (https://www.gisaid.org), despite high vaccination rates in adults and/or high prevalence of prior infection^28^.

We demonstrate evasion of neutralising antibodies by a B.1.617.2 live virus with sera from convalescent patients, as well as sera from individuals vaccinated with two different vaccines, one based on an adenovirus vector (ChAdOx-1), and the other mRNA based (BNT162b2). Our findings on reduced susceptibility of B.1.617.2 to vaccine elicited sera are similar to other reports^29,30^, including the lower GMT following two doses of ChAdOx-1 compared to BNT162b2^29^. Although we did not map the mutations responsible, previous work with shows that L452R and T478K in the spike RBD are likely to have contributed^10^, as well as spike NTD mutations. The importance of NTD in both cell entry efficiency^24,31^ as well as antibody evasion is increasingly recognised^32,33^ and further work is needed to map specific determinants in the B.1.617.2 NTD.

We also report ChAdOx-1 vaccine breakthrough infections in health care workers at three Delhi hospitals. These infections were predominantly B.1.617.2, with a mix of other lineages including B.1.1.7, reflecting prevalence in community infections. We estimated the relative VE of ChAdOx-1 vaccination in our HCW analysis against B.1.617.2 versus other lineages, finding an increased odds of symptomatic infection and disease with B.1.617.2 compared to non-B.1.617.2 following two doses. These data indicate reduced VE against B.1.617.2 and support an immune evasion advantage for B.1.617.2.

It is important to consider that increased infectivity at mucosal surfaces and cell-cell fusion and spread^34^ may also facilitate ‘evasion’ from antibodies^35^. Indeed, our work also shows that that B.1.617.2 had a fitness advantage compared to B.1.1.7 across physiologically relevant systems including HAE and 3D airway organoids^20^ where cell free and cell-cell infection are likely to be occurring together. These data support the notion of higher infectiousness of B.1.617.2, either due to higher viral burden or higher particle infectivity, resulting in higher probability of person-to-person transmission. We noted that B.1.617.2 live virus particles contained a higher proportion of cleaved spike compared to B.1.1.7, and postulated that this is involved in the mechanism of increased infectivity. Consistent with this hypothesis, we observed that PV particles bearing B.1.617.2 spike demonstrated significantly enhanced entry into a range of target cells.

The B.1.617.1 variant was detected before B.1.617.2 in India, and the reasons for B.1.617.2 out-competing B.1.617.1 are unknown. We report that B.1.617.2 has a replication advantage in lung cells compared to B.1.617.1, and that this is reflected in a PV entry advantage driven by spike. Given our data showing that B.1.617.2 and B.1.617.1 spikes confer similar sensitivities to sera from vaccinees, superior fitness is a parsimonious explanation for the growth advantage of B.1.617.2 over B.1.617.1.

Virus infectivity and fusogenicity mediated by the PBCS is a key determinant of pathogenicity and transmissibility^19,36^ and there are indications that giant cells/syncytia formation are associated with fatal disease^37^. Spike cleavage and stability of cleaved spike are likely therefore to be critical parameters for future SARS-CoV-2 variants of concern. B.1.617.2 spike demonstrated similar kinetics of syncytia formation as compared to B.1.1.7, likely attributable to P681R. We show that vaccine-elicited sera can inhibit syncytia formation, and that this blockade of cell-cell fusion is compromised for B.1.617.2, potentially also permitting virus to pass from cell to cell and thereby evading neutralising antibodies generated following vaccination.

The REGN-COV2 dual monoclonal antibody therapy containing casirivimab and imedevimab was shown to improve survival for non-B.1.617.2 infections ^38^. Reduced efficacy for imedevimab against B.1.617.2 shown here could translate to compromised clinical efficacy. Moreover, it could lead to possible selection of escape variants where there is immune compromise and chronic SARS-CoV-2 infection with B.1.6 1 7.2^39^. Further work to explore these possibilities is urgently needed.

Although protection against infection with B.1.351 (the variant with least sensitivity to neutralising antibodies) has been demonstrated for at least three vaccines^13,40–42^, progression to severe disease and death has been low. Therefore, at population scale, extensive vaccination will likely protect against moderate to severe disease due to B.1.617.2. Indeed data from the UK already demonstrate low incidence of severe disease in vaccinees (PHE technical report 17). However, our data on vaccine breakthrough and reduced vaccine effectiveness against symptomatic B.1.617.2 infection are of concern given that hospitals frequently treat individuals who may have suboptimal immune responses to vaccination due to comorbidity. Such patients could be at risk for severe disease following infection from HCW and indeed we document here a ‘super-spreading’ event involving infection vaccinated HCWs. Therefore strategies to boost vaccine responses against variants are warranted and attention to infection control procedures is needed in the post vaccine era.

## Methods

### Serum samples and ethical approval

Ethical approval for study of vaccine elicited antibodies in sera from vaccinees was obtained from the East of England – Cambridge Central Research Ethics Committee Cambridge (REC ref: 17/EE/0025). Use of convalescent sera had ethical approval from South Central Berkshire B Research Ethics Committee (REC ref: 20/SC/0206; IRAS 283805). Testing and sequencing of positive samples for genomic surveillance is part of Indian government mandated responsibilities of National Centres for Disease Control and CSIR-IGIB for public health purposes. Research related to these activities was approved by The Institutional Human Ethics Committee (NCDC/**2020/NERC/14 and** CSIR-IGIB/IHEC/2020-21/01)

Studies involving testing and sequencing of positive samples from health care workers were reviewed and approved by The Institutional Human Ethics Committees of NCDC and CSIR-IGIB(NCDC/**2020/NERC/14 and** CSIR-IGIB/IHEC/2020-21/01)

### Sequencing Quality Control and Phylogenetic Analysis

Three sets of fasta concensus sequences were obtained from three separate Hospitals in Delhi, India. Initially, all sequences were concatenated into a multi-fasta, according to hospital, and then aligned to reference strain MN908947.3 (Wuhan-Hu-1) with mafft v4.475 ^43^ using the --keeplength --addfragments options. Following this, all sequences were passed through Nextclade v0.15 (https://clades.nextstrain.org/) to determine the number of gap regions. This was noted and all sequences were assigned a lineage with Pangolin v3.1.5^44^ and pangoLEARN (dated 15^th^ June 2021). Sequences that could not be assigned a lineage were discarded. After assigning lineages, all sequences with more than 5% N-regions were also excluded.

Phylogenies were inferred using maximum-likelihood in IQTREE v2.1.4^45^ using a GTR+R6 model with 1000 rapid bootstraps. The inferred phylogenies were annotated in R v4.1.0 using ggtree v3.0.2^46^ and rooted on the SARS-CoV-2 reference sequence (MN908947.3). Nodes were arranged in descending order and lineages were annotated on the phylogeny as coloured tips, alongside a heatmap defining the number of ChAdOx-1 vaccines received from each patient.

### Structural Analyses

The PyMOL Molecular Graphics System v.2.4.0 (https://github.com/schrodinger/pymol-open-source/releases) was used to map the location of the mutations defining the Delta lineage (B.1.617.2) onto closed-conformation spike protein - PDB: 6ZGE^47^.

### Statistical Analyses

#### Vaccine breakthrough infections in Health care workers

Descriptive analyses of demographic and clinical data are presented as median and interquartile range (IQR) or mean and standard deviation (SD) when continuous and as frequency and proportion (%) when categorical. The difference in continuous and categorical data were tested using Wilcoxon rank sum or T-test and Chi-square test respectively. The association between Ct value and SARS-CoV-2 variant was examined using linear regression. Variants as the dependent variable were categorized into two groups: B.1.617.2 variant and non-B.1.617.2 variants. The following covariates were included in the model irrespective of confounding: age, sex, hospital and interval between symptom onset and nasal swab PCR testing.

#### Vaccine effectiveness

To estimate vaccine effectiveness (VE) for the B.1.617.2 variant relative to non-B.1.617.2 variants, we adopted a recently described approach^27^. This method is based on the premise that if the vaccine is equally effective against B.1.617.2 and non-B.1.617.2 variants, a similar proportion of cases with either variant would be expected in both vaccinated and unvaccinated cases. This approach overcomes the issue of higher background prevalence of one variant over the other. We determined the proportion of cases with the B.1.617.2 variant relative to all other circulating variants by vaccination status. We then used a logistic regression to estimate the odds ratio of testing positive with B.1.617.2 in vaccinated compared to unvaccinated individuals. The final regression model was adjusted for age as a continuous variable, sex and hospital as categorical variables. Model sensitivity and robustness to inclusion of these covariates was tested by an iterative process of sequentially adding the covariates to the model and examining the impact on the ORs and confidence internals until the final model was constructed (**Extended Data Table 4**). The R-square measure, as proposed by McFadden^48^, was used to test the fit of different specifications of the same model regression. This is was done by sequential addition of the variables adjusted for including age, sex and hospital until the final model was constructed. In addition, the absolute difference in Bayesian Information Criterion (BIC) was estimated. The McFadden R^2^ measure of final model fitness was 0.11 indicating reasonable model fit. The addition of age, gender and hospital in the final regression model improved the measured fitness. However, the absolute difference in BIC was 13.34 between the full model and the model excluding the adjusting variable, providing strong support for the parsimonious model. The fully adjusted model was nonetheless used as the final model as the sensitivity analyses (**Extended Data Table 4**) showed robustness to the addition of the covariates.

#### Neutralisation titre analyses

The neutralisation by vaccine-elicited antibodies after the two doses of the BNT162b2 and Chad-Ox-1 vaccine was determined by infections in the presence of serial dilutions of sera as described below. The ID50 within groups were summarised as a geometric mean titre (GMT) and statistical comparison between groups were made with Mann-Whitney or Wilcoxon ranked sign test. Statistical analyses were done using Stata v13 and Prism v9.

### Pseudotype virus experiments

#### Cells

HEK 293T CRL-3216, Hela-ACE-2 (Gift from James Voss), Vero CCL-81 were maintained in Dulbecco’s Modified Eagle Medium (DMEM) supplemented with 10% fetal calf serum (FCS), 100 U/ml penicillin, and 100mg/ml streptomycin. All cells were regularly tested and are mycoplasma free. H1299 cells were a kind gift from Sam Cook. Calu-3 cells were a kind gift from Paul Lehner, A549 A2T2^49^ cells were a kind gift from Massimo Palmerini. Vero E6 Ace2/TMPRSS2 cells were a kind gift from Emma Thomson.

#### Pseudotype virus preparation for testing against vaccine elicited antibodies and cell entry

Plasmids encoding the spike protein of SARS-CoV-2 D614 with a C terminal 19 amino acid deletion with D614G were used. Mutations were introduced using Quickchange Lightning Site-Directed Mutagenesis kit (Agilent) following the manufacturer’s instructions. B.1.1.7 S expressing plasmid preparation was described previously, but in brief was generated by step wise mutagenesis. Viral vectors were prepared by transfection of 293T cells by using Fugene HD transfection reagent (Promega). 293T cells were transfected with a mixture of 11ul of Fugene HD, 1μg of pCDNAΔ19 spike-HA, 1ug of p8.91 HIV-1 gag-pol expression vector and 1.5μg of pCSFLW (expressing the firefly luciferase reporter gene with the HIV-1 packaging signal). Viral supernatant was collected at 48 and 72h after transfection, filtered through 0.45um filter and stored at −80°C as previously described. Infectivity was measured by luciferase detection in target 293T cells transfected with TMPRSS2 and ACE2.

#### Standardisation of virus input by SYBR Green-based product-enhanced PCR assay (SG-PERT)

The reverse transcriptase activity of virus preparations was determined by qPCR using a SYBR Green-based product-enhanced PCR assay (SG-PERT) as previously described^50^. Briefly, 10-fold dilutions of virus supernatant were lysed in a 1:1 ratio in a 2x lysis solution (made up of 40% glycerol v/v 0.25% Triton X-100 v/v 100mM KCl, RNase inhibitor 0.8 U/ml, TrisHCL 100mM, buffered to pH7.4) for 10 minutes at room temperature.

12μl of each sample lysate was added to thirteen 13μl of a SYBR Green master mix (containing 0.5μM of MS2-RNA Fwd and Rev primers, 3.5pmol/ml of MS2-RNA, and 0.125U/μl of Ribolock RNAse inhibitor and cycled in a QuantStudio. Relative amounts of reverse transcriptase activity were determined as the rate of transcription of bacteriophage MS2 RNA, with absolute RT activity calculated by comparing the relative amounts of RT to an RT standard of known activity.

### Viral isolate comparison between B.1.617.1 and B.1.617.2

#### Cell Culture

VeroE6/TMPRSS2 cells [an African green monkey (*Chlorocebus sabaeus*) kidney cell line; JCRB1819]^51^ were maintained in Dulbecco’s modified Eagle’s medium (low glucose) (Wako, Cat# 041-29775) containing 10% FCS, G418 (1 mg/ml; Nacalai Tesque, Cat# G8168-10ML) and 1% antibiotics (penicillin and streptomycin; PS).

Calu-3 cells (a human lung epithelial cell line; ATCC HTB-55) were maintained in Minimum essential medium Eagle (Sigma-Aldrich, Cat# M4655-500ML) containing 10% FCS and 1% PS.

#### SARS-Co V-2 B.1.617.1 vs B.1.617.2 experiment

Two viral isolates belonging to the B.1.617 lineage, B.1.617.1 (GISAID ID: EPI_ISL_2378733) and B.1.617.2 (GISAID ID: EPI_ISL_2378732) were isolated from SARS-CoV-2-positive individuals in Japan. Briefly, 100 μl of the nasopharyngeal swab obtained from SARS-CoV-2-positive individuals were inoculated into VeroE6/TMPRSS2 cells in the biosafety level 3 laboratory. After the incubation at 37°C for 15 minutes, a maintenance medium supplemented [Eagle’s minimum essential medium (FUJIFILM Wako Pure Chemical Corporation, Cat# 056-08385) including 2% FCS and 1% PS] was added, and the cells were cultured at 37°C under 5% CO_2_. The cytopathic effect (CPE) was confirmed under an inverted microscope (Nikon), and the viral load of the culture supernatant in which CPE was observed was confirmed by real-time RT-PCR. To determine viral genome sequences, RNA was extracted from the culture supernatant using QIAamp viral RNA mini kit (Qiagen, Qiagen, Cat# 52906). cDNA library was prepared by using NEB Next Ultra RNA Library Prep Kit for Illumina (New England Biolab, Cat# E7530) and whole genome sequencing was performed by Miseq (Illumina).

To prepare the working virus, 100 μl of the seed virus was inoculated into VeroE6/TMPRSS2 cells (5,000,000 cells in a T-75 flask). At one hour after infection, the culture medium was replaced with Dulbecco’s modified Eagle’s medium (low glucose) (Wako, Cat# 041-29775) containing 2% FBS and 1% PS; at 2-3 days postinfection, the culture medium was harvested and centrifuged, and the supernatants were collected as the working virus.

The titer of the prepared working virus was measured as 50% tissue culture infectious dose (TCID_50_). Briefly, one day prior to infection, VeroE6/TMPRSS2 cells (10,000 cells/well) were seeded into a 96-well plate. Serially diluted virus stocks were inoculated to the cells and incubated at 37°C for 3 days. The cells were observed under microscopy to judge the CPE appearance. The value of TCID_50_/ml was calculated with the Reed-Muench method^52^.

One day prior to infection, 20, 000 Calu-3 cells were seeded into a 96-well plate. SARS-CoV-2 (200 TCID_50_) was inoculated and incubated at 37°C for 1 h. The infected cells were washed, and 180 μl of culture medium was added. The culture supernatant (10 μl) was harvested at indicated time points and used for real-time RT-PCR to quantify the viral RNA copy number.

#### Real-Time RT-PCR

Real-time RT-PCR was performed as previously described^53,54^. Briefly, 5 μl of culture supernatant was mixed with 5 μl of 2 × RNA lysis buffer [2% Triton X-100, 50 mM KCl, 100 mM Tris-HCl (pH 7.4), 40% glycerol, 0.8 U/μl recombinant RNase inhibitor (Takara, Cat# 2313B)] and incubated at room temperature for 10 min. RNase-free water (90 μl) was added, and the diluted sample (2.5 μl) was used as the template for real-time RT-PCR performed according to the manufacturer’s protocol using the One Step TB Green PrimeScript PLUS RT-PCR kit (Takara, Cat# RR096A) and the following primers: Forward *N*, 5’-AGC CTC TTC TCG TTC CTC ATC AC-3’; and Reverse *N*, 5’-CCG CCA TTG CCA GCC ATT C-3’. The copy number of viral RNA was standardized with a SARS-CoV-2 direct detection RT-qPCR kit (Takara, Cat# RC300A). The fluorescent signal was acquired using a QuantStudio 3 Real-Time PCR system (Thermo Fisher Scientific), a CFX Connect RealTime PCR Detection system (Bio-Rad) or a 7500 Real Time PCR System (Applied Biosystems).

#### Virus growth kinetics in HAE cells

Primary nasal human airway epithelial (HAE) cells at air-liquid interface (ALI) were purchased from Epithelix and the basal MucilAir medium (Epithelix) was changed every 2-3 days for maintenance of HAE cells. All dilution of viruses, wash steps and harvests were carried out with OptiPRO SFM (Life Technologies) containing 2X GlutaMAX (Gibco). All wash and harvest steps were performed by addition of 200ul SFM to the apical surface and incubation for 10 mins at 37°C before removing SFM. To infect, basal medium was replaced, the apical surface of the HAE cells washed once with SFM to remove mucus before addition of virus to triplicate wells. Cells were infected at a multiplicity of 1e4 genomes copies of virus per cell based on E gene qRT-PCR. Inoculum was incubated for 1 h at 37°C before removing, washing the apical surface twice and the second wash taken as harvest for 0 hpi. A single apical wash was performed to harvest virus at 24, 48 and 71 hr timepoints. Isolates used were B.1.617.2 isolate #60 hCoV-19/England/SHEF-10E8F3B/2021 (EPI_ISL_1731019), B.1.617.2 isolate #285 hCoV-19/England/PHEC-3098A2/2021 (EPI_ISL_2741645) and B.1.1.7 isolate #7540 SMH2008017540 (confirmed B.1.1.7 inhouse but not yet available on GISAID).

| Subtype | Name | Sequencing | 19 | 77 | 142 | 156-158 | 222 | 452 | 478 | 614 | 681 | 950 |
| --- | --- | --- | --- | --- | --- | --- | --- | --- | --- | --- | --- | --- |
| Wuhan | Wuhan | - | T | K | G | WT | A | L | T | D | P | D |
| B.1.617.2 | 60 | VAT2 | R | R | D | del | V | R | K | G | R | N |
| B.1.617.2 | 285 | VAT2 | R |  | D | del | V | R | K | G | R | N |

#### Titration of outputs from HAE infections

For determining genome copies in the virus inputs and in the supernatant harvested from HAE cells, RNA was extracted using QIAsymphony DSP Virus/Pathogen Mini Kit on the QIAsymphony instrument (Qiagen). qRT-PCR was then performed using AgPath RT-PCR (Life Technologies) kit on a QuantStudio(TM) 7 Flex System with the primers for SARS-CoV-2 E gene used in Corman et al., (2020). A standard curve was also generated using dilutions viral RNA of known copy number to allow quantification of E gene copies in the samples from Ct values. E gene copies per ml of original virus supernatant were then calculated.

For measuring infectious virus in harvests from HAE cells, plaque assays were performed by performing serial log dilutions of supernatant in DMEM, 1% NEAA and 1% P/S and inoculating onto PBS-washed Vero cells, incubating for 1 hr at 37°C, removing inoculum and overlaying with 1× MEM, 0.2% w/v BSA, 0.16% w/v NaHCO3, 10□mM HEPES, 2mM L-Glutamine, 1× P/S, 0.6% w/v agarose. Plates were incubated for 3 d at 37□°C before overlay was removed and cells were stained for 1□h at room temperature in crystal violet solution.

#### Lung organoid infection by replication competent SARS-CoV-2 isolates

Airway epithelial organoids were prepared as previously reported.^20^ For viral infection primary organoids were passaged and incubated with SARS-CoV-2 in suspension at a multiplicity of infection (MOI) of 1 for 2 hours. Subsequently, the infected organoids were washed twice with PBS to remove the viral particles. Washed organoids were plated in 20 μl Matrigel domes, cultured in organoid medium and harvested at different timepoints.

Cells were lysed 24 and 48h post-infection and total RNA isolated. cDNA was synthesized and qPCR was used to determine copies of nucleoprotein gene in samples. Standard curve was prepared using SARS-CoV-2 Positive Control plasmid containing full nucleocapsid protein (N gene) (NEB) and used to quantify copies of N gene in organoid samples. 18S ribosomal RNA was used as a housekeeping gene to normalize sample-to-sample variation.

#### Western blotting

Cells were lysed and supernatants collected 18 hours post transfection. Purified virions were prepared by harvesting supernatants and passing through a 0.45 μm filter. Clarified supernatants were then loaded onto a thin layer of 8.4% optiprep density gradient medium (Sigma-Aldrich) and placed in a TLA55 rotor (Beckman Coulter) for ultracentrifugation for 2 hours at 20,000 rpm. The pellet was then resuspended for western blotting. Cells were lysed with cell lysis buffer (Cell signalling), treated with Benzonase Nuclease (70664 Millipore) and boiled for 5 min. Samples were then run on 4%–12% Bis Tris gels and transferred onto nitrocellulose or PVDF membranes using an iBlot or semidry (Life Technologies and Biorad, respectively).

Membranes were blocked for 1 hour in 5% non-fat milk in PBS + 0.1% Tween-20 (PBST) at room temperature with agitation, incubated in primary antibody (anti-SARS-CoV-2 Spike, which detects the S2 subunit of SARS-CoV-2 S (Invitrogen, PA1-41165), anti-GAPDH (proteintech) or anti-p24 (NIBSC)) diluted in 5% non-fat milk in PBST for 2 hours at 4°C with agitation, washed four times in PBST for 5 minutes at room temperature with agitation and incubated in secondary antibodies anti-rabbit HRP (1:10000, Invitrogen 31462), anti-bactin HRP (1:5000; sc-47778) diluted in 5% non-fat milk in PBST for 1 hour with agitation at room temperature. Membranes were washed four times in PBST for 5 minutes at room temperature and imaged directly using a ChemiDoc MP imaging system (Bio-Rad).

#### Virus infection for virion western blotting

Vero-hACE2-TMPRS22 cells were infected with MOI of 1 and incubated for 48 hours. Supernatant was cleared by 5 min spin at 300xg and then precipitated with 10% PEG6000 (4h at RT). Pellets were resuspended directly in Laemmli buffer with 1mM DTT, then treated with Benzonase Nuclease(70664 Millipore) and sonicated prior loading for gel electrophoresis

### Serum pseudotype neutralisation assay

Spike pseudotype assays have been shown to have similar characteristics as neutralisation testing using fully infectious wild type SARS-CoV-2^55^.Virus neutralisation assays were performed on 293T cell transiently transfected with ACE2 and TMPRSS2 using SARS-CoV-2 spike pseudotyped virus expressing luciferase^56^. Pseudotyped virus was incubated with serial dilution of heat inactivated human serum samples or convalescent plasma in duplicate for 1h at 37°C. Virus and cell only controls were also included. Then, freshly trypsinized 293T ACE2/TMPRSS2 expressing cells were added to each well. Following 48h incubation in a 5% CO_2_ environment at 37°C, the luminescence was measured using Steady-Glo Luciferase assay system (Promega).

#### Neutralization Assays for convalescent plasma

Convalescent sera from healthcare workers at St. Mary’s Hospital at least 21 days since PCR-confirmed SARS-CoV-2 infection were collected in May 2020 as part of the REACT2 study. Convalescent human serum samples were inactivated at 56°C for 30 min and replicate serial 2-fold dilutions (n=12) were mixed with an equal volume of SARs-CoV-2 (100 TCID50; total volume 100 μL) at 37°C for 1□h. Vero-hACE2 TMPRSS2 cells were subsequently infected with serial-fold dilutions of each sample for 3 days at 37°C. Virus neutralisation was quantified via crystal violet staining and scoring for cytopathic effect (CPE). Each-run included 1/5 dilutions of each test sample in the absence of virus to ensure virus-induced CPE in each titration. Back-titrations of SARs-CoV-2 infectivity were performed to demonstrate infection with ~100 TCID50 in each well.

### Vaccinee Serum neutralization, live virus assays

Vero-Ace2-TMPRSS2 cells were seeded at a cell density of 2×10e4/well in 96w plate 24h before infection. Serum was titrated starting at a final 1:10 dilution with WT (SARS-CoV-2/human/Liverpool/REMRQ0001/2020), B.1.1.7 or B.1.617.2 virus isolates being added at MOI 0.01. The mixture was incubated 1h prior adding to cells. The plates were fixed with 8% PFA 72h post-infection and stained with Coomassie blue for 20 minutes. The plates were washed in water and dried for 2h. 1% SDS was added to wells and staining intensity was measured using FLUOstar Omega (BMG Labtech). Percentage cell survival was determined by comparing intensity of staining to an uninfected wells. A non-linear sigmoidal 4PL model (Graphpad Prism 9.1.2) was used to determine the ID50 for each serum.

### VSV pseudovirus generation for monoclonal antibody assays

Replication defective VSV pseudovirus expressing SARS-CoV-2 spike proteins corresponding to the different VOC were generated as previously described with some modifications^57^. Lenti-X 293T cells (Takara, 632180) were seeded in 10-cm^2^ dishes at a density of 5e6 cells per dish and the following day transfected with 10 μg of WT or B.1.617.2 spike expression plasmid with TransIT-Lenti (Mirus, 6600) according to the manufacturer’s instructions. One day post-transfection, cells were infected with VSV-luc (VSV-G) with an MOI of 3 for 1 h, rinsed three times with PBS containing Ca2+/Mg2+, then incubated for an additional 24 h in complete media at 37°C. The cell supernatant was clarified by centrifugation, filtered (0.45 um), aliquoted, and frozen at −80°C.

#### Pseudotyped virus neutralization assay for mAb

Vero E6 expressing TMPRSS2 or not were grown in DMEM supplemented with 10% FBS and seeded into white 96 well plates (PerkinElmer, 6005688) at a density of 20 thousand cells per well. The next day, mAbs were serially diluted in pre-warmed complete media, mixed with WT or B.1.617.2 pseudoviruses and incubated for 1 h at 37°C in round bottom polypropylene plates. Media from cells was aspirated and 50 μl of virus-mAb complexes were added to cells and then incubated for 1 h at 37°C. An additional 100 μL of pre-warmed complete media was then added on top of complexes and cells incubated for an additional 16-24 h. Conditions were tested in duplicate wells on each plate and at least six wells per plate contained untreated infected cells (defining the 0% of neutralization, “MAX RLU” value) and infected cells in the presence of S2E12 and S2X259 at 25 μg/ml each (defining the 100% of neutralization, “MIN RLU” value). Virus-mAb-containing media was then aspirated from cells and 50 μL of a 1:2 dilution of SteadyLite Plus (Perkin Elmer, 6066759) in PBS with Ca^++^ and Mg^++^ was added to cells. Plates were incubated for 15 min at room temperature and then were analysed on the Synergy-H1 (Biotek). Average of Relative light units (RLUs) of untreated infected wells (MAX RLUave) was subtracted by the average of MIN RLU (MIN RLU_ave_) and used to normalize percentage of neutralization of individual RLU values of experimental data according to the following formula: (1-(RLU_x_ - MIN RLU_ave_) / (MAX RLU_ave_ – MIN RLU_ave_)) x 100. Data were analyzed and visualized with Prism (Version 9.1.0). IC50 values were calculated from the interpolated value from the log(inhibitor) versus response, using variable slope (four parameters) nonlinear regression with an upper constraint of ≤100, and a lower constrain equal to 0. Each neutralization assay was conducted on two independent experiments, i.e., biological replicates, where each biological replicate contains a technical duplicate. IC50 values across biological replicates are presented as arithmetic mean ± standard deviation. The loss or gain of neutralization potency across spike variants was calculated by dividing the variant IC50 by the WT IC50 within each biological replicate, and then visualized as arithmetic mean ± standard deviation.

#### Plasmids for split GFP system to measure cell-cell fusion

pQCXIP□BSR□GFP11 and pQCXIP□GFP1□10 were from Yutaka Hata^58^ Addgene plasmid #68716; http://n2t.net/addgene:68716; RRID:Addgene_68716 and Addgene plasmid #68715; http://n2t.net/addgene:68715; RRID:Addgene_68715)

#### Generation of GFP1□10 or GFP11 lentiviral particles

Lenti viral particles were generated by co-transfection of Vero cells with pQCXIP□BSR□GFP11 or pQCXIP□GFP1□10 as previously described^59^. Supernatant containing virus particles was harvested after 48 and 72 hours, 0.45 μm filtered, and used to infect 293T or Vero cells to generate stable cell lines. 293T and Vero cells were transduced to stably express GFP1□10 or GFP11 respectively and were selected with 2 μg/ml puromycin.

#### Cell-cell fusion assay

Cell-cell fusion assay was carried out as previously described^59,60^ but using a Split-GFP system. Briefly, Vero GFP1-10 and Vero-GFP11 cells were seeded at 80% confluence in a 1:1 ration in 24 multiwell plate the day before. Cells. were co-transfected with 0.5 μg of spike expression plasmids in pCDNA3 using Fugene 6 and following the manufacturer’s instructions (Promega). Cell-cell fusion was measured using an Incucyte and determined as the proportion of green area to total phase area. Data were then analysed using Incucyte software analysis. Graphs were generated using Prism 8 software.

## Supporting information

EDTABLE1

EDTABLE2

EDTABLE3

EDTABLE3

## Data availability

All fasta consensus sequences files used in this analysis are available from https://gisaid.org or from https://github.com/Steven-Kemp/hospital_india/tree/main/consensus_fasta. Code for the Bayesian modelling analysis is available at: https://github.com/ImperialCollegeLondon/delta_modelling

## Author contributions

Conceived study: AA, PR, SAK, DC, SB, SF, SM, RKG, DAC, LP, JB, DP, DC. Designed study and experiments: BM, RKG, JB, NG, LCJ, GP, KS, IATM. Performed experiments: BM, DAC, RD, IATMF, CS-F, CS, JB, LCG, JB, JZ, NG, GBM. Interpreted data: RKG, SAK, AA, SS, JB, RP, PC, PD, KP, VSR, SS, DC, TP, OC, KS, GP, LCJ, WB, SF, SB, DAC, BM, RD, IATMF, PR, JB, KGCS, SM, CW, TM, SB, LP, DC, CS, CS-F and S.F.

## Acknowledgments

We would like to thank the Department of Biotechnology, NCDC, RKG is supported by a Wellcome Trust Senior Fellowship in Clinical Science (WT108082AIA). This study was supported by the Cambridge NIHRB Biomedical Research Centre. We would also like to thank Ankur Mutreja. We would like to thank Thushan de Silva for the Delta isolate and Kimia Kimelian. SAK is supported by the Bill and Melinda Gates Foundation via PANGEA grant: OPP1175094. I.A.T.M.F. is funded by a SANTHE award (DEL-15-006). We would like to thank Paul Lehner for Calu-3 cells. We would like to thank Clare Lloyd and Sejal Saglani for providing the primary airway epithelial cultures, and James Voss for HeLa ACE2. We thank the Geno2pheno UK consortium. The authors acknowledge support from the G2P-UK National Virology consortium funded by MRC/UKRI (grant ref: MR/W005611/1).This study was also supported by The Rosetrees Trust and the Geno2pheno UK consortium. SF acknowledges the EPSRC (EP/V002910/1). KS is supported by AMED Research Program on Emerging and Re-emerging Infectious Diseases (20fk0108270 and 20fk0108413), JST SICORP (JPMJSC20U1 and JPMJSC21U5) and JST CREST (JPMJCR20H4).

## INSACOG CONSORTIUM MEMBERS

**NIBMG:** Saumitra Das, Arindam Maitra, Sreedhar Chinnaswamy, Nidhan Kumar Biswas;

**ILS:** Ajay Parida, Sunil K Raghav, Punit Prasad;

**InSTEM/ NCBS**: Apurva Sarin, Satyajit Mayor, Uma Ramakrishnan, Dasaradhi Palakodeti, Aswin Sai Narain Seshasayee;

**CDFD**: K Thangaraj, Murali Dharan Bashyam, Ashwin Dalal;

**NCCS**: Manoj Bhat, Yogesh Shouche, Ajay Pillai;

**IGIB:** Anurag Agarwal, Sridhar Sivasubbu, Vinod Scaria;

**NIV:** Priya Abraham, Potdar Varsha Atul, Sarah S Cherian;

**NIMHANS:** Anita Sudhir Desai, Chitra Pattabiraman, M. V. Manjunatha, Reeta S Mani, Gautam Arunachal Udupi;

**NCDC:** Sujeet Singh, Himanshu Chauhan, Partha Rakshit, Tanzin Dikid;

**CCMB: Vinay Nandicoori**, Karthik Bharadwaj Tallapaka, Divya Tej Sowpati

## INSACOG collaborating divisions and clinical partners

Biotechnology division, NCDC: Hema Gogia, Hemlata Lall, Kalaiarasan Ponnusamy, Kaptan Verma, Mahesh S Dhar, Manoj K Singh, Meena Datta, Namita Soni, Namonarayan Meena, Partha Rakshit, Preeti Madan, Priyanka Singh, Radhakrishnan V. S, Ramesh Sharma, Rajeev Sharma, Robin Marwal, Sandhya Kabra, Sattender Kumar, Swati Kumari, Uma Sharma, Urmila Chaudhary Integrated Disease Surveillance Program (IDSP), NCDC, Central and State IDSP units Centre of Epidemiology, NCDC

Sri Ganga Ram Hospital, Rajinder Nagar, New Delhi: Dept of Clinical Microbiology & Immunology and Director Medical Hospital Administration, Chand Wattal, J K Oberoi, Neeraj Goel, Reena Raveendran, S. Datta

Northern Railway Central Hospital, Basant Lane, New Delhi: Meenakshi Agarwal Indraprastha Apollo Hospitals, New Delhi: Administration and Microbiology department, Raju Vaishya

## The Genotype to Phenotype Japan (G2P-Japan) Consortium members

**The Institute of Medical Science, The University of Tokyo**: Jumpei Ito, Izumi Kimura, Keiya Uriu, Yusuke Kosugi, Mai Suganami, Akiko Oide, Miyabishara Yokoyama, Mika Chiba

**Hiroshima University**: Ryoko Kawabata, Nanami Morizako

**Tokyo Metropolitan Institute of Public Health**: Kenji Sadamasu, Hiroyuki Asakura, Mami Nagashima, Kazuhisa Yoshimura

**University of Miyazaki**: Akatsuki Saito, Erika P Butlertanaka, Yuri L Tanaka

**Kumamoto University**: Terumasa Ikeda, Chihiro Motozono, Hesham Nasser, Ryo Shimizu, Yue Yuan, Kazuko Kitazato, Haruyo Hasebe

**Tokai University**: So Nakagawa, Jiaqi Wu, Miyoko Takahashi

**Hokkaido University**: Takasuke Fukuhara, Kenta Shimizu, Kana Tsushima, Haruko Kubo

**Kyoto University**: Kotaro Shirakawa, Yasuhiro Kazuma, Ryosuke Nomura, Yoshihito Horisawa, Akifumi Takaori-Kondo

**National Institute of Infectious Diseases**: Kenzo Tokunaga, Seiya Ozono

## The CITIID-NIHR BioResource COVID-19 Collaboration

### Principal Investigators

Ravindra K Gupta^1,2,3^, Stephen Baker^2, 3^, Gordon Dougan^2, 3^, Christoph Hess^2,3,28,29^, Nathalie Kingston^22, 12^, Paul J. Lehner^2,20,3^, Paul A. Lyons^2, 3^, Nicholas J. Matheson^2, 3^, Willem H. Owehand^22^, Caroline Saunders^21^, Charlotte Summers^3,26,27,30^, James E.D. Thaventhiran^2, 3, 24^, Mark Toshner^3, 26, 27^, Michael P. Weekes^2,20^, Patrick Maxwell^22,30^, Ashley Shaw^30^

### CRF and Volunteer Research Nurses

Ashlea Bucke^21^, Jo Calder^21^, Laura Canna^21^, Jason Domingo^21^, Anne Elmer^21^, Stewart Fuller^21^, Julie Harris^43^, Sarah Hewitt^21^, Jane Kennet^21^, Sherly Jose^21^, Jenny Kourampa^21^, Anne Meadows^21^, Criona O’Brien^43^, Jane Price^21^, Cherry Publico^21^, Rebecca Rastall^21^, Carla Ribeiro^21^, Jane Rowlands^21^, Valentina Ruffolo^21^, Hugo Tordesillas^21^,

### Sample Logistics

Ben Bullman^2^, Benjamin J. Dunmore^3^, Stuart Fawke^32^, Stefan Gräf^3,22,12^, Josh Hodgson^3^, Christopher Huang^3^, Kelvin Hunter^2 3^, Emma Jones^31^, Ekaterina Legchenko^3^, Cecilia Matara^3^, Jennifer Martin^3^, Federica Mescia^2, 3^, Ciara O’Donnell^3^, Linda Pointon^3^, Nicole Pond^2, 3^, Joy Shih^3^, Rachel Sutcliffe^3^, Tobias Tilly^3^, Carmen Treacy^3^, Zhen Tong^3^, Jennifer Wood^3^, Marta Wylot^38^,

### Sample Processing and Data Acquisition

Laura Bergamaschi^2, 3^, Ariana Betancourt^2, 3^, Georgie Bower^2, 3^, Chiara Cossetti^2, 3^, Aloka De Sa^3^, Madeline Epping^2, 3^, Stuart Fawke^32^, Nick Gleadall^22^, Richard Grenfell^33^, Andrew Hinch^2,3^, Oisin Huhn^34^, Sarah Jackson^3^, Isobel Jarvis^3^, Ben Krishna^3^, Daniel Lewis^3^, Joe Marsden^3^, Francesca Nice^41^, Georgina Okecha^3^, Ommar Omarjee^3^, Marianne Perera^3^, Martin Potts^3^, Nathan Richoz^3^, Veronika Romashova^2,3^, Natalia Savinykh Yarkoni^3^, Rahul Sharma^3^, Luca Stefanucci^22^, Jonathan Stephens^22^, Mateusz Strezlecki^33^, Lori Turner^2, 3^,

### Clinical Data Collection

Eckart M.D.D. De Bie^3^, Katherine Bunclark^3^, Masa Josipovic^42^, Michael Mackay^3^, Federica Mescia^2,3^, Alice Michael^27^, Sabrina Rossi^37^, Mayurun Selvan^3^, Sarah Spencer^15^, Cissy Yong^37^

### Royal Papworth Hospital ICU

Ali Ansaripour^27^, Alice Michael^27^, Lucy Mwaura^27^, Caroline Patterson^27^, Gary Polwarth^27^

### Addenbrooke’s Hospital ICU

Petra Polgarova^30^, Giovanni di Stefano^30^

### Cambridge and Peterborough Foundation Trust

Codie Fahey^36^, Rachel Michel^36^

### ANPC and Centre for Molecular Medicine and Innovative Therapeutics

Sze-How Bong^23^, Jerome D. Coudert^35^, Elaine Holmes^39^

### NIHR BioResource

John Allison^22,12^, Helen Butcher^12,40^, Daniela Caputo^12,40^, Debbie Clapham-Riley^12,40^, Eleanor Dewhurst^12,40^, Anita Furlong^12,40^, Barbara Graves^12,40^, Jennifer Gray^12,40^, Tasmin Ivers^12,40^, Mary Kasanicki^12,30^, Emma Le Gresley^12,40^, Rachel Linger^12,40^, Sarah Meloy^12,40^, Francesca Muldoon^12,40^, Nigel Ovington^22,12^, Sofia Papadia^12,40^, Isabel Phelan^12,40^, Hannah Stark^12,40^, Kathleen E Stirrups^22,12^, Paul Townsend^22,12^, Neil Walker^22,12^, Jennifer Webster^12,40^, Ingrid Scholtes^40^, Sabine Hein^40^, Rebecca King^40^

^1^University College London, Infection & Immunity, London, UK

^2^Cambridge Institute of Therapeutic Immunology & Infectious Disease, Cambridge, UK.

^3^Department of Medicine, University of Cambridge, Cambridge, UK.

^4^Humabs Biomed SA, a subsidiary of Vir Biotechnology, 6500 Bellinzona, Switzerland.

^5^Department of Biochemistry, University of Washington, Seattle, WA 98195, USA

^6^Vir Biotechnology, San Francisco, CA 94158, USA.

^7^Clinic of Internal Medicine and Infectious Diseases, Clinica Luganese Moncucco, 6900 Lugano, Switzerland

^8^Division of Infectious Diseases, Luigi Sacco Hospital, University of Milan, Milan, Italy

^9^ The CITIID-NIHR BioResource COVID-19 Collaboration, see appendix 1 for author list

^10^ NIHR Cambridge Clinical Research Facility, Cambridge, UK.

^11^ NIHR Bioresource, Cambridge, UK

^12^ University of Kent, Canturbury, UK

^13^Department of Clinical Biochemistry and Immunology, Addenbrookes Hospital, UK

^14^ Laboratorio de Inmunologia, S-Cuautitlán, UNAM, Mexico

^16^ Institute of Biodiversity, University of Glasgow, Glasgow, UK

^17^Department of Haematology, University of Cambridge, Cambridge CB2 0QQ, UK

^18^University of KwaZulu Natal, Durban, South Africa

^19^Africa Health Research Institute, Durban, South Africa

^20^Dept of Infectious Diseases, Cambridge University Hospitals NHS Trust, Cambridge UK.

^21^Cambridge Clinical Research Centre, NIHR Clinical Research Facility, Cambridge University Hospitals NHS Foundation Trust, Addenbrooke’s Hospital, Cambridge CB2 0QQ, UK

^22^University of Cambridge, Cambridge Biomedical Campus, Cambridge CB2 0QQ, UK

^23^Australian National Phenome Centre, Murdoch University, Murdoch, Western Australia WA 6150, Australia

^24^MRC Toxicology Unit, School of Biological Sciences, University of Cambridge, Cambridge CB2 1QR, UK

^25^R&D Department, Hycult Biotech, 5405 PD Uden, The Netherlands

^26^Heart and Lung Research Institute, Cambridge Biomedical Campus, Cambridge CB2 0QQ, UK

^27^Royal Papworth Hospital NHS Foundation Trust, Cambridge Biomedical Campus, Cambridge CB2 0QQ, UK

^28^Department of Biomedicine, University and University Hospital Basel, 4031Basel, Switzerland

^29^Botnar Research Centre for Child Health (BRCCH) University Basel & ETH Zurich, 4058 Basel, Switzerland

^30^Addenbrooke’s Hospital, Cambridge CB2 0QQ, UK

^31^Department of Veterinary Medicine, Madingley Road, Cambridge, CB3 0ES, UK

^32^Cambridge Institute for Medical Research, Cambridge Biomedical Campus, Cambridge CB2 0XY, UK

^33^Cancer Research UK, Cambridge Institute, University of Cambridge CB2 0RE, UK

^34^Department of Obstetrics & Gynaecology, The Rosie Maternity Hospital, Robinson Way, Cambridge CB2 0SW, UK

^35^Centre for Molecular Medicine and Innovative Therapeutics, Health Futures Institute, Murdoch University, Perth, WA, Australia

^36^Cambridge and Peterborough Foundation Trust, Fulbourn Hospital, Fulbourn, Cambridge, UK

^37^Department of Surgery, Addenbrooke’s Hospital, Cambridge CB2 0QQ, UK

^38^Department of Biochemistry, University of Cambridge, Cambridge, CB2 1QW, UK

^39^Centre of Computational and Systems Medicine, Health Futures Institute, Murdoch University, Harry Perkins Building, Perth, WA 6150, Australia

^40^Department of Public Health and Primary Care, School of Clinical Medicine, University of Cambridge, Cambridge Biomedical Campus, Cambridge, UK

---

**Extended Data Figure 1.**
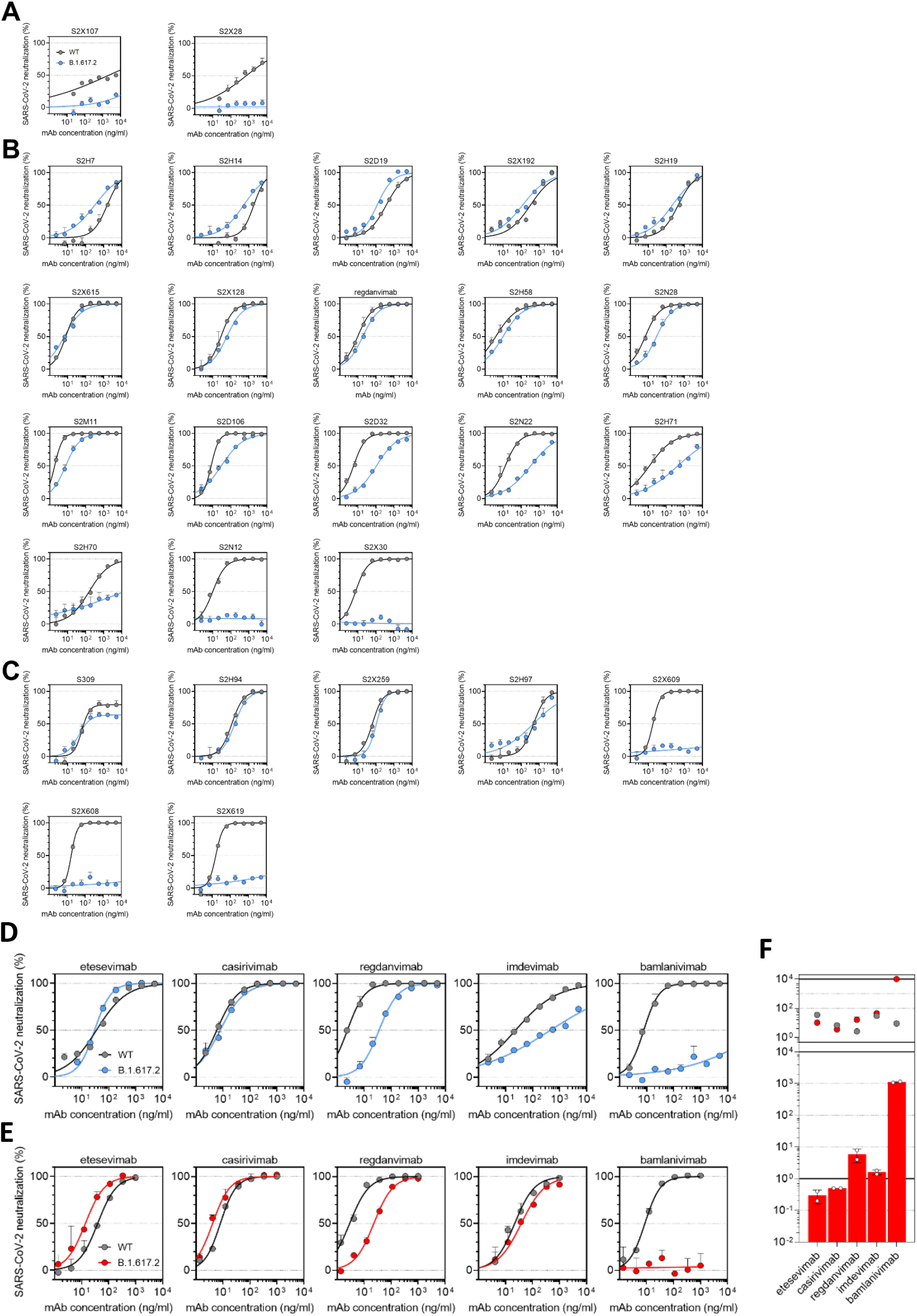
Neutralisation by a panel of NTD- and RBD-specific mAbs against WT and B.1.617.2 mutant SARS-CoV-2 pseudotyped viruses. **a-c**, Neutralisation of WT (Wuhan-1 D614, black) and B.1.617.2 mutant (blue) pseudotyped SARS-CoV-2-VSV by 27 mAbs targeting NTD (A, n=3), RBM (B, n=18) and non-RBM (C, n=7). Shown is one representative experiment out of 2 independent experiments. **d-e**, Neutralisation of WT D614 (black) and B.1.617.2 mutant (blue/red) pseudotyped SARS-CoV-2-VSV by 5 clinical-stage mAbs using Vero E6 cells expressing TMPRSS2 (d) or not (e). Shown is one representative experiment out of 2 independent experiments. **f,** Neutralisation shown as mean IC50 values (upper panel) and average fold change of B.1.617.2 relative to WT (lower panel) of 5 clinical mAbs tested in 2 independent experiments using Vero E6 cells not expressing TMPRSS2 (related to panel e).

**Extended Data Figure 2.**
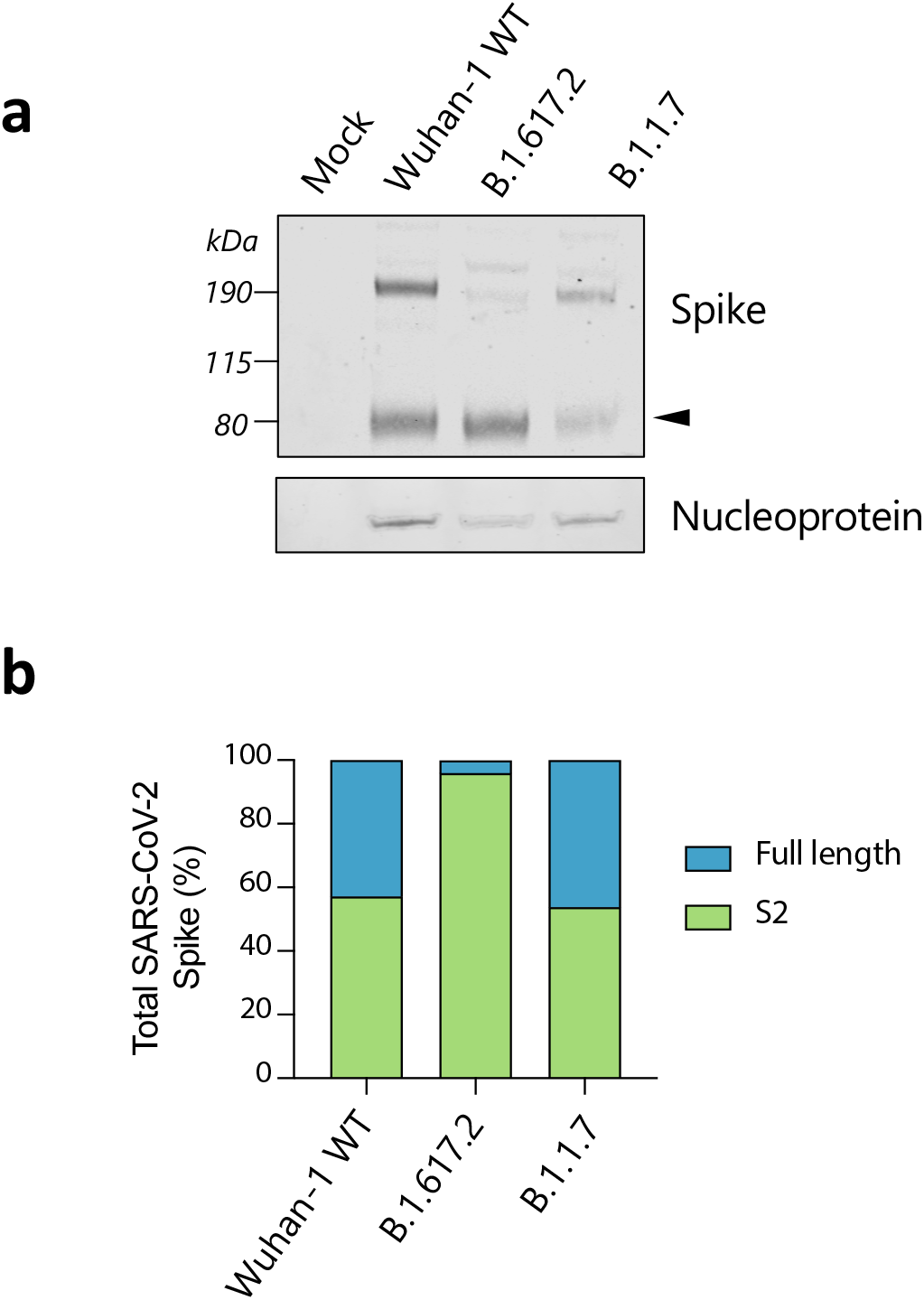
Spike cleavage in B.1.617.2 virions compared to B.1.1.7. **a.** Representative western blot analysis of spike and nucleoprotein present in SARS-CoV-2 particles from the indicated viruses produced in Vero-hACE2-TMPRS22 cells 48 hours post infection. The arrowhead identifies the S2 subunit. **b**. Quantification of cleaved and fulllength spike of the indicated viruses.

**Extended Data Figure 3:**
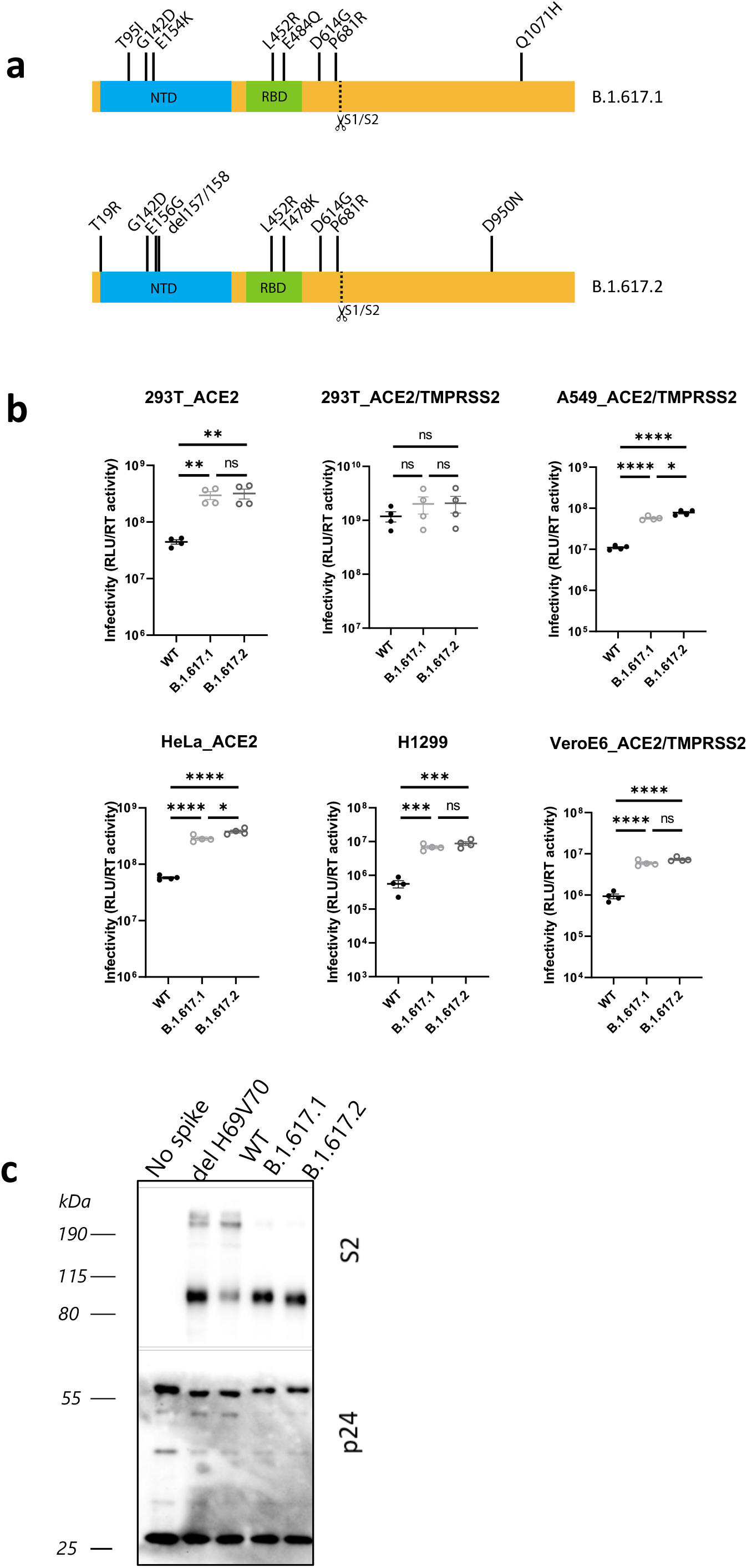
B.1.617.2 spike confers increased cell entry. **a.** diagram showing mutations present in spike plasmids used for cell entry PV experiments **b.** Single round infectivity on different cell targets by spike B.1.617.1 and B.1.617.1 versus WT (Wuhan-1 D614G) PV produced in 293T cells. Data are representative of three independent experiments. Statistics were performed using unpaired Student t test. **c.** Western blotting of supernatants from transfected 293T probing for S2 and p24 in PV and showing no spike control.

**Extended Data Figure 4.**
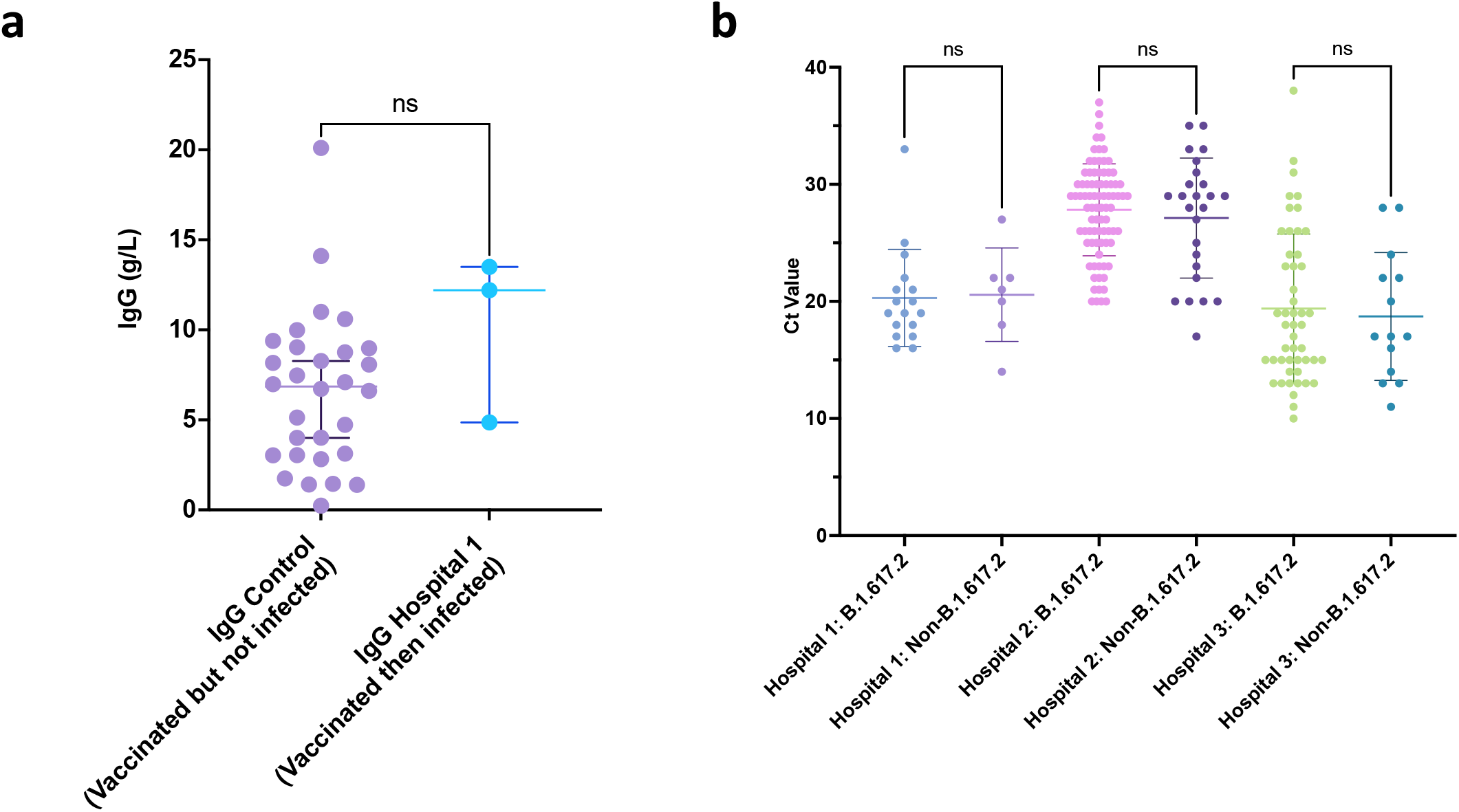
Breakthrough SARS-CoV-2 infections amongst vaccinated health care workers (HCW) **a.** Comparison of IgG antibody titres between a control group of vaccinated individuals receiving two doses of ChadOx-1 who have not been infected with SARS-CoV-2, with vaccinated healthcare workers who had received two doses and subsequently tested positive for SARS-CoV-2. **b.** Ct values in nose/throat swabs from HCW testing positive by hospital. Bars represent Mean and 95% CI. Ct values were compared using the Student t test.

## References

1 Volz, E. et al. Assessing transmissibility of SARS-CoV-2 lineage B.1.1.7 in England. Nature, doi:10.1038/s41586-021-03470-x (2021).

2 Collier, D. A. et al. SARS-CoV-2 B.1.1.7 sensitivity to mRNA vaccine-elicited, convalescent and monoclonal antibodies. Nature 593, 136–141, doi:10.1101/2021.01.19.21249840 (2021).

3 Cherian, S. et al. Convergent evolution of SARS-CoV-2 spike mutations, L452R, E484Q and P681R, in the second wave of COVID-19 in Maharashtra, India. bioRxiv, 2021.2004.2022.440932, doi:10.1101/2021.04.22.440932 (2021).

4 Hoffmann, M. et al. SARS-CoV-2 variant B.1.617 is resistant to Bamlanivimab and evades antibodies induced by infection and vaccination. bioRxiv, 2021.2005.2004.442663, doi:10.1101/2021.05.04.442663 (2021).

5 Ferreira, I. et al. SARS-CoV-2 B.1.617 emergence and sensitivity to vaccine-elicited antibodies. bioRxiv, 2021.2005.2008.443253, doi:10.1101/2021.05.08.443253 (2021).

6 Yadav, P. D. et al. Neutralization of variant under investigation B.1.617 with sera of BBV152 vaccinees. bioRxiv, 2021.2004.2023.441101, doi:10.1101/2021.04.23.441101 (2021).

7 Deng, X. et al. Transmission, infectivity, and antibody neutralization of an emerging SARS-CoV-2 variant in California carrying a L452R spike protein mutation. medRxiv, doi:10.1101/2021.03.07.21252647 (2021).

8 Ferreira, I. et al. SARS-CoV-2 B.1.617 mutations L452 and E484Q are not synergistic for antibody evasion. The Journal of infectious diseases, doi:10.1093/infdis/jiab368 (2021).

9 McCallum, M. et al. SARS-CoV-2 immune evasion by variant B.1.427/B.1.429. bioRxiv, 2021.2003.2031.437925, doi:10.1101/2021.03.31.437925 (2021).

10 Motozono, C. et al. An emerging SARS-CoV-2 mutant evading cellular immunity and increasing viral infectivity. bioRxiv, 2021.2004.2002.438288, doi:10.1101/2021.04.02.438288 (2021).

11 Hadfield, J. et al. Nextstrain: real-time tracking of pathogen evolution. Bioinformatics 34, 4121–4123, doi:10.1093/bioinformatics/bty407 (2018).

12 JHU. CORONA VIRUS RESOURCE CENTRE, <https://coronavirus.jhu.edu/map.html> (2021).

13 Madhi, S. A. et al. Efficacy of the ChAdOx1 nCoV-19 Covid-19 Vaccine against the B.1.351 Variant. N Engl J Med, doi:10.1056/NEJMoa2102214 (2021).

14 Kustin, T. et al. Evidence for increased breakthrough rates of SARS-CoV-2 variants of concern in BNT162b2 mRNA vaccinated individuals. medRxiv, 2021.2004.2006.21254882, doi:10.1101/2021.04.06.21254882 (2021).

15 Collier, D. A. et al. Age-related immune response heterogeneity to SARS-CoV-2 vaccine BNT162b2. Nature, doi:10.1038/s41586-021-03739-1 (2021).

16 Van Oekelen, O. et al. Highly variable SARS-CoV-2 spike antibody responses to two doses of COVID-19 RNA vaccination in patients with multiple myeloma. Cancer Cell, doi:10.1016/j.ccell.2021.06.014 (2021).

17 Chen, P. et al. SARS-CoV-2 Neutralizing Antibody LY-CoV555 in Outpatients with Covid-19. New England Journal of Medicine 384, 229–237, doi:10.1056/NEJMoa2029849 (2020).

18 Weinreich, D. M. et al. REGN-COV2, a Neutralizing Antibody Cocktail, in Outpatients with Covid-19. New England Journal of Medicine 384, 238–251, doi:10.1056/NEJMoa2035002 (2020).

19 Peacock, T. P. et al. The furin cleavage site in the SARS-CoV-2 spike protein is required for transmission in ferrets. Nat Microbiol, doi:10.1038/s41564-021-00908-w (2021).

20 Youk, J. et al. Three-Dimensional Human Alveolar Stem Cell Culture Models Reveal Infection Response to SARS-CoV-2. Cell Stem Cell 27, 905–919 e910, doi:10.1016/j.stem.2020.10.004 (2020).

21 Hoffmann, M. et al. SARS-CoV-2 Cell Entry Depends on ACE2 and TMPRSS2 and Is Blocked by a Clinically Proven Protease Inhibitor. Cell 181, 271–280 e278, doi:10.1016/j.cell.2020.02.052 (2020).

22 Papa, G. et al. Furin cleavage of SARS-CoV-2 Spike promotes but is not essential for infection and cell-cell fusion. PLoS pathogens 17, e1009246, doi:10.1371/journal.ppat.1009246 (2021).

23 Cattin-Ortolá, J. et al. Sequences in the cytoplasmic tail of SARS-CoV-2 Spike facilitate expression at the cell surface and syncytia formation. bioRxiv, 2020.2010.2012.335562, doi:10.1101/2020.10.12.335562 (2021).

24 Meng, B. et al. Recurrent emergence and transmission of a SARS-CoV-2 spike deletion H69/V70 and role in Alpha Variant B.1.1.7. Cell reports, doi:https://doi.org/10.1016/j.celrep.2021.109292 (2021).

25 Winstone, H. et al. The Polybasic Cleavage Site in SARS-CoV-2 Spike Modulates Viral Sensitivity to Type I Interferon and IFITM2. Journal of virology 95, e02422–02420, doi:10.1128/jvi.02422-20 (2021).

26 Bhoyar, R. C. et al. High throughput detection and genetic epidemiology of SARS-CoV-2 using COVIDSeq next-generation sequencing. PloS one 16, e0247115, doi: 10.1371/journal.pone.0247115 (2021).

27 Lopez Bernal, J. et al. Effectiveness of Covid-19 Vaccines against the B.1.617.2 (Delta) Variant. New England Journal of Medicine, doi:10.1056/NEJMoa2108891 (2021).

28 Bernal, J. L. et al. Effectiveness of COVID-19 vaccines against the B.1.617.2 variant. medRxiv, 2021.2005.2022.21257658, doi:10.1101/2021.05.22.21257658 (2021).

29 Wall, E. C. et al. Neutralising antibody activity against SARS-CoV-2 VOCs B.1.617.2 and B.1.351 by BNT162b2 vaccination. Lancet, doi:10.1016/S0140-6736(21)01290-3 (2021).

30 Planas, D. et al. Reduced sensitivity of infectious SARS-CoV-2 variant B.1.617.2 to monoclonal antibodies and sera from convalescent and vaccinated individuals. bioRxiv, 2021.2005.2026.445838, doi:10.1101/2021.05.26.445838 (2021).

31 Lempp, F. A. et al. Membrane lectins enhance SARS-CoV-2 infection and influence the neutralizing activity of different classes of antibodies. bioRxiv, 2021.2004.2003.438258, doi:10.1101/2021.04.03.438258 (2021).

32 McCarthy, K. R. et al. Natural deletions in the SARS-CoV-2 spike glycoprotein drive antibody escape. Science (2020).

33 McCallum, M. et al. N-terminal domain antigenic mapping reveals a site of vulnerability for SARS-CoV-2. Cell 184, 2332–2347 e2316, doi:10.1016/j.cell.2021.03.028 (2021).

34 Zeng, C. et al. SARS-CoV-2 Spreads through Cell-to-Cell Transmission. bioRxiv, 2021.2006.2001.446579, doi:10.1101/2021.06.01.446579 (2021).

35 Jackson, L. et al. SARS-CoV-2 cell-to-celI spread occurs rapidly and is insensitive to antibody neutralization. bioRxiv, 2021.2006.2001.446516, doi:10.1101/2021.06.01.446516 (2021).

36 Johnson, B. A. et al. Loss of furin cleavage site attenuates SARS-CoV-2 pathogenesis. Nature 591, 293–299, doi:10.1038/s41586-021-03237-4 (2021).

37 Braga, L. et al. Drugs that inhibit TMEM16 proteins block SARS-CoV-2 Spike-induced syncytia. Nature, doi:10.1038/s41586-021-03491-6 (2021).

38 Group, R. C. et al. Casirivimab and imdevimab in patients admitted to hospital with COVID-19 (RECOVERY): a randomised, controlled, open-label, platform trial. medRxiv, 2021.2006.2015.21258542, doi:10.1101/2021.06.15.21258542 (2021).

39 Kemp, S. A. et al. SARS-CoV-2 evolution during treatment of chronic infection. Nature 592, 277–282, doi:10.1038/s41586-021-03291-y (2021).

40 Shinde, V. et al. Efficacy of NVX-CoV2373 Covid-19 Vaccine against the B.1.351 Variant. New England Journal of Medicine, doi:10.1056/NEJMoa2103055 (2021).

41 Abu-Raddad, L. J., Chemaitelly, H. & Butt, A. A. Effectiveness of the BNT162b2 Covid-19 Vaccine against the B.1.1.7 and B.1.351 Variants. New England Journal of Medicine, doi:10.1056/NEJMc2104974 (2021).

42 Sadoff, J. et al. Safety and Efficacy of Single-Dose Ad26.COV2.S Vaccine against Covid-19. New England Journal of Medicine, doi:10.1056/NEJMoa2101544 (2021).

43 Katoh, K. & Standley, D. M. MAFFT multiple sequence alignment software version 7: improvements in performance and usability. Mol Biol Evol 30, 772–780, doi:10.1093/molbev/mst010 (2013).

44 Rambaut, A. et al. A dynamic nomenclature proposal for SARS-CoV-2 lineages to assist genomic epidemiology. Nat Microbiol 5, 1403–1407, doi: 10.1038/s41564-020-0770-5 (2020).

45 Minh, B. Q. et al. IQ-TREE 2: New models and efficient methods for phylogenetic inference in the genomic era. bioRxiv, 849372, doi:10.1101/849372 (2019).

46 Yu, G., Smith, D. K., Zhu, H., Guan, Y. & Lam, T. T. Y. ggtree: an R package for visualization and annotation of phylogenetic trees with their covariates and other associated data. Methods in Ecology and Evolution 8, 28–36 (2017).

47 Wrobel, A. G. et al. SARS-CoV-2 and bat RaTG13 spike glycoprotein structures inform on virus evolution and furin-cleavage effects. Nat Struct Mol Biol 27, 763–767, doi:10.1038/s41594-020-0468-7 (2020).

48 McFadden, D. Conditional logit analysis of qualitative choice behavior. 105–142 (Academic Press, 1974).

49 Rihn, S. J. et al. A plasmid DNA-launched SARS-CoV-2 reverse genetics system and coronavirus toolkit for COVID-19 research. PLoS Biol 19, e3001091, doi:10.1371/journal.pbio.3001091 (2021).

50 Vermeire, J. et al. Quantification of reverse transcriptase activity by real-time PCR as a fast and accurate method for titration of HIV, lenti- and retroviral vectors. PloS one 7, e50859–e50859, doi:10.1371/journal.pone.0050859 (2012).

51 Matsuyama, S. et al. Enhanced isolation of SARS-CoV-2 by TMPRSS2-expressing cells. Proc Natl Acad Sci U S A 117, 7001–7003, doi:10.1073/pnas.2002589117 (2020).

52 Reed, L. J. & Muench, H. A Simple Method of Estimating Fifty Percent Endpoints. Am J Hygiene 27, 493–497 (1938).

53 Motozono, C. et al. SARS-CoV-2 spike L452R variant evades cellular immunity and increases infectivity. Cell Host Microbe, doi:10.1016/j.chom.2021.06.006 (2021).

54 Shema Mugisha, C. et al. A simplified quantitative real-time PCR assay for monitoring SARS-CoV-2 growth in cell culture. mSphere 5,doi:10.1128/mSphere.00658-20 (2020).

55 Schmidt, F. et al. Measuring SARS-CoV-2 neutralizing antibody activity using pseudotyped and chimeric viruses. 2020.2006.2008.140871, doi:10.1101/2020.06.08.140871 %J bioRxiv (2020).

56 Mlcochova, P. et al. Combined point of care nucleic acid and antibody testing for SARS-CoV-2 following emergence of D614G Spike Variant. Cell Rep Med, 100099, doi:10.1016/j.xcrm.2020.100099 (2020).

57 Ou, X. et al. Characterization of spike glycoprotein of SARS-CoV-2 on virus entry and its immune cross-reactivity with SARS-CoV. Nat Commun 11, 1620, doi:10.1038/s41467-020-15562-9 (2020).

58 Kodaka, M. et al. A new cell-based assay to evaluate myogenesis in mouse myoblast C2C12 cells. Experimental cell research 336, 171–181 (2015).

59 Papa, G. et al. Furin cleavage of SARS-CoV-2 Spike promotes but is not essential for infection and cell-cell fusion. PLoS Pathogens 17, e1009246 (2021).

60 Buchrieser, J. et al. Syncytia formation by SARS-CoV-2-infected cells. The EMBO journal 39, e106267 (2020).

