## Supplementary material for "SARS-CoV-2 B.1.617.2 Delta variant replication, sensitivity to neutralising antibodies and vaccine breakthrough": EDTABLE1

**Extended Data Table 1** Demographic information for individuals undergoing two dose SARS-CoV-2 vaccination with ChAdOx-1 or BNT162b2

| | ChAdOx-1 (N=33) | BNT162b2 (N=32) | P. value |
|---|---|---|---|
| Female (%) | 18 (54.5) | 13 (40.6) | 0.38[a] |
| Median Age *Years* (IQR) | 67 (64 - 71) | 71 (46 -83) | 0.74[b] |
| Median time *in Days* since dose 2 (IQR) | 31 (21 -38) | 27 (24 -29) | 0.15[b] |
| Serum Geometric Mean Titre *GMT* for delta variant (95% CI) | 654 (313 -1365) | 3372 (1856 - 6128) | 0.0006[b] |
| Serum Geometric Mean Titre *GMT* for WT (95% CI) | 2625 (1492 - 4618) | 7393 (3893 - 14041) | 0.0030[b] |

[a]Chi-square test, [b] Mann-Whitney test
