## Supplementary material for "SARS-CoV-2 B.1.617.2 Delta variant replication, sensitivity to neutralising antibodies and vaccine breakthrough": EDTABLE2

| mAb | Domain/site | IC50 WT (ng/ml) | IC50 B.1.617.2 (ng/ml) | VH usage (% id.) | Source (DSO) | ACE2 blocking | Ref. |
|---|---|---|---|---|---|---|---|
| **S2X107** | **NTD** | 2611.00 | 10000.00 | 4-38-2 (97) | Sympt. (75) | Neg. | McCallum et al. |
| **S2X28** | **NTD** | 1121.30 | 10000.00 | 3-30 (97.9) | Sympt. (48) | Neg. | McCallum et al. |
| **S2X333** | **NTD** | 217.95 | 35016.00 | 3-33 (96.5) | Sympt. (125) | Neg. | McCallum et al. |
| **S2H7** | **RBM** | 1227.25 | 347.05 | 3-66 (98.3) | Sympt. (17) | Weak | Thomson et al. |
| **S2H14** | **RBM** | 1666.00 | 566.40 | 3-15 (100) | Sympt. (17) | Weak | Piccoli et al.; Thomson et al. |
| **S2D19** | **RBM** | 369.55 | 129.95 | 4-31 (99.7) | Hosp. (49) | Moderate | Thomson et al. |
| **S2X192** | **RBM** | 423.00 | 246.15 | 1-69 (96.9) | Sympt. (75) | Weak | Thomson et al. |
| **S2H19** | **RBM** | 549.25 | 382.45 | 3-15 (98.6) | Sympt. (45) | Weak | Thomson et al. |
| **S2E12** | **RBM** | 2.09 | 1.72 | 1-58 (97.6) | Hosp. (51) | Strong | Thomson et al.; Tortorici et al. |
| **S2X615** | **RBM** | 9.80 | 14.59 | 3-11 (94.8) | Sympt. (271) | Strong | Collier et al. |
| **S2X128** | **RBM** | 36.82 | 65.97 | 1-69-2 (97.6) | Sympt. (75) | Strong | Thomson et al. |
| **S2D8** | **RBM** | 8.44 | 16.78 | 3-23 (96.5) | Hosp. (49) | Strong | Thomson et al. |
| **S2H58** | **RBM** | 5.21 | 12.55 | 1-2 (97.9) | Sympt. (45) | Strong | Thomson et al. |
| **S2N28** | **RBM** | 10.05 | 36.95 | 3-30 (97.2) | Hosp. (51) | Strong | Thomson et al. |
| **S2M11** | **RBM** | 2.07 | 8.32 | 1-2 (96.5) | Hosp. (46) | Weak | Thomson et al.; Tortorici et al. |
| **S2D106** | **RBM** | 8.0 | 34.6 | 1-69 (97.2) | Hosp. (98) | Strong | Thomson et al. |
| **S2D32** | **RBM** | 4.6 | 104.6 | 3-49 (98.3) | Hosp. (49) | Strong | Thomson et al. |
| **S2N22** | **RBM** | 16.8 | 543.4 | 3-23 (96.5) | Hosp. (51) | Strong | Thomson et al. |
| **S2D97** | **RBM** | 10.0 | 332.9 | 2-5 (96.9) | Hosp. (98) | Weak | Thomson et al. |
| **S2H71** | **RBM** | 18.2 | 622.8 | 2-5 (99) | Sympt. (45) | Moderate | Thomson et al. |
| **S2H70** | **RBM** | 194.9 | 10000.0 | 1-2 (99) | Sympt. (45) | Weak | Thomson et al. |
| **S2X58** | **RBM** | 14.4 | 10000.0 | 1-46 (99) | Sympt. (48) | Strong | Thomson et al. |
| **S2N12** | **RBM** | 10.6 | 10000.0 | 4-39 (97.6) | Hosp. (51) | Strong | Thomson et al. |
| **S2X30** | **RBM** | 10.0 | 10000.0 | 1-69 (97.9) | Sympt. (48) | Strong | Thomson et al. |
| **etesevimab** | **RBM** | 28.7 | 23.0 / 10.3* | 3-66 (99.7) | Sympt. (?) | Strong | R. Shi et al. Nature 2020 |
| **casirivimab** | **RBM** | 6.0 | 7.2 / 3.5* | 3-30 (98.6) | Immunized mice | Strong | J. Hansen et al. Science 2020 |
| **regdanvimab** | **RBM** | 2.2 | 29.2 / 16.3* | 2-70 ( ?) | Sympt. (?) | Strong | |
| **imdevimab** | **RBM** | 31.9 | 1607.2 / 45.4* | 3-11 (98.6) | Sympt. (?) | Strong | J. Hansen et al.Science 2020 |
| **bamlanivimab** | **RBM** | 7.8 | 10000 / 10000* | 1-69 (99.7) | Sympt. (?) | Strong | Jones et al.Sci Transl Med 2021 |
| **S309** | **non-RBM** | 63.9 | 46.4 | 1-18 (97.2) | SARS-CoV | Weak | Pinto et al. |
| **S2X35** | **non-RBM** | 181.2 | 224.6 | 1-18 (98.6) | Sympt. (48) | Strong | Piccoli et al. |
| **S2H94** | **non-RBM** | 134.1 | 182.9 | 3-23 (93.4) | Sympt. (81) | Strong | Thomson et al. |
| **S2X259** | **non-RBM** | 74.2 | 101.1 | 1-69 (94.1) | Sympt. (75) | Moderate | Tortorici et al (BioRxiv 2021) |
| **S2H97** | **non-RBM** | 599.3 | 1260.4 | 5-51 (98.3) | Sympt. (81) | Weak | Collier et al.; Starr et al (BioRxiv 21) |
| **S2X609** | **non-RBM** | 19.7 | 10000.0 | 1-69 (93.8) | Sympt. (271) | Strong | Collier et al. |
| **S2X608** | **non-RBM** | 18.9 | 10000.0 | 1-33 (93.2) | Sympt. (271) | Strong | Collier et al. |
| **S2X619** | **non-RBM** | 18.2 | 10000.0 | 1-69 (92.7) | Sympt. (271) | Strong | Collier et al. |
| **S2X305** | **non-RBM** | 11.8 | 9522.5 | 1-2 (95.1) | Sympt. (125) | Strong | Collier et al. |

DSO, days after symptom onset. Sympt., symptomatic. Hosp., hospitalized. SARS-CoV, infected with SARS-CoV virus. *clinical-stage mAbs tested with Vero E6 cells not expressing TMPRSS2

**Extended Data Table 2:** Monoclonal antibodies used in neutralisation assays against pseudotyped virus bearing spike from WT (Wuhan-1 D614) or B.1.617.2.
