## Supplementary material for "SARS-CoV-2 B.1.617.2 Delta variant replication, sensitivity to neutralising antibodies and vaccine breakthrough": EDTABLE3

**Extended Data Table 3: Data on SARS-CoV-2 infections in three hospitals with near universal staff vaccination during first half of 2021.**

| | B.1.617.2 (N= 112) | Non-B.1.617.2 (N=20) | P value |
|---|---|---|---|
| Median age *years* (IQR) | 36.5 (27.0-49.5) | 32.5 (27.5-44.0) | 0.56[a] |
| Female % | 51.8 (58) | 50.0 (10) | 0.88[b] |
| Hospital % | | | |
| 1 | 9.8 (11) | 15.0 (3) | 0.15[b] |
| 2 | 53.6 (60) | 30.0 (6) | |
| 3 | 36.6 (41) | 55.0 (11) | |
| Median Ct value (IQR) | 22.5 (16.4-28.6)[c] | 19.8 (17.3-22.8) | 0.48 |
| Number of vaccines doses %[†] | | | |
| 0 | 10.8 (12) | 35.0 (7) | 0.005[b] |
| 1 | 20.7 (23) | 30.0 (6) | |
| 2 | 68.5 (76) | 35.0 (7) | |
| Hospitalised %* | | | |
| No | 95.5 (64) | 93.3 (14) | 0.72[b] |
| Yes | 4.5 (3) | 6.7 (1) | |
| Anti-Spike IgG GMT (95% CI) | 15.5 (4.6-52.9)[d] | 29.5 (0.0-2.4x$10^{6}$)[e] | 0.69[a] |
| Median Symptom duration *days* | 1.5 (1.0-3.0)[f] | 1.0 (1.0-2.0)[g] | 0.66[a] |

[a] Wilcoxon rank-sum test. [b]Chi square test. [c]111 of 112 available. [d]11 of 112. [e]2 of 20. [f]63 of 112. [g] 12 of 20. [†]Vaccine status missing for 1 of 132. *Hospitalisation data is unavailable from Hospital 1. IQR- inrerquartile range, GMT- geometric mean titre. CI- confidence interval.
