## Supplementary material for "SARS-CoV-2 B.1.617.2 Delta variant replication, sensitivity to neutralising antibodies and vaccine breakthrough": EDTABLE3

**Extended Data Table 4**: **Relative ChAdOx-1 vaccine effectiveness against B1.617.2 v non- B1.617.2:** Odds ratios for detection of B.1.617.2 relative to non-B.1.617.2 in vaccinated compared to unvaccinated individuals in multi-variable logistic regression. Bottom table below shows sensitivity of model to iterative addition of covariates. OR; odds ratio aOR; Adjusted odds ratio.

| | B1.617.2 | Non-B.1.617.2 | B1.617.2: Non-B.1.617.2 | OR (95% CI) | P value | aOR (95% CI) | P value |
|---|---|---|---|---|---|---|---|
| Unvaccinated | 12 | 7 | 1.71 | - | | - | |
| Vaccinated | | | | | | | |
| Dose 1 | 23 | 6 | 3.83 | 2.24 (0.61-8.16) | 0.22 | 2.18 (0.53-9.01) | 0.28 |
| Dose 2 | 76 | 7 | 10.86 | 6.33 (1.89-21.27) | 0.003 | 5.45 (1.39-21.4) | 0.015 |
| Dose 1 and 2 | 99 | 13 | 7.62 | 4.44 (1.48-13.30) | 0.008 | 3.81 (1.11-13.03) | 0.03 |

| Model includes covariates | OR for B.1.617.2 vs non-B.1.617.2 (95% CI) | P value | OR for age (95% CI) | P value | OR for sex (95% CI) | P value | OR for hospital (95% CI) | P value |
|---|---|---|---|---|---|---|---|---|
| Dose 1 and 2 | 4.44 (1.48-13.30) | 0.008 | | | | | | |
| +age | 4.23 (1.34-13.31) | 0.014 | 1.01 (0.97-1.05) | 0.78 | | | | |
| +sex | 4.43 (1.48-13.29) | 0.008 | | | 0.96 (0.36-2.57) | 0.93 | | |
| +hospital<br>Hospital 1<br>Hospital 2<br>Hospital 3 | 4.64 (1.45-14.80) | 0.01 | | | | | <br>Baseline<br>3.71 (0.77-17.94)<br>1.54 (0.34-6.96) | <br>-<br>0.10<br>0.58 |
| +age +sex | 4.14 (1.29-13.28) | 0.017 | 1.01 (0.96-1.05) | 0.74 | 0.89 (0.31-2.60) | 0.84 | | |
| +age +sex +hospital<br>Hospital 1<br>Hospital 2<br>Hospital 3 | 3.81 (1.11- 13.03) | 0.03 | 1.03 (0.98-1.09) | 0.22 | 1.54 (0.48-4.97) | 0.47 | <br><br>Baseline<br>8.69 (1.19-63.37)<br>2.27 (0.45-11.53) | <br><br>-<br>0.03<br>0.32 |
